# Discovery of Abundant Nano-scale Lymphatic-like Vessels in Brains

**DOI:** 10.64898/2025.12.30.697095

**Authors:** Shiju Gu, Hongquan Dong, Hao Chen, Jiang Yu, Lei Liu, Jinwu Yan, Huizhe Wang, Zhiyong Jiang, Wanqian Huang, Wei Wang, Steven H. Liang, Can Zhang, Shiqian Shen, Chongzhao Ran

**Author notes:** These authors contribute equally to this work (Hongquan Dong, Hao Chen).

## Abstract

As one of the most metabolically active organs, the brain requires an exceptionally efficient system for waste clearance to sustain its high metabolic demands. However, whether such a system exists—and, if so, what structural features enable its efficiency—remains incompletely understood. More than a decade ago, the “glymphatic system” was proposed to describe neurofluid transport through cerebrospinal fluid (CSF) and perivascular spaces (PVS), in conjunction with dural and meningeal lymphatic pathways. Nevertheless, it remains unresolved whether neurofluid transport is organized as a structured, vessel-like flow network. Moreover, in stark contrast to the dense network of blood capillaries in the brain, only a sparse population of lymphatic vessels has been identified, raising doubts as to whether the currently recognized lymphatic architecture alone can support efficient metabolic waste clearance.

By combining expansion microscopy with CRANAD-3, a pan–β-amyloid fluorescent probe, we discovered abundant nanoscale lymphatic-like vessels (NLVs) within the brain parenchyma of both mice and humans. The majority of these structures have diameters below 1,000 nm and exhibit moderate positivity for multiple lymphatic markers, including LYVE-1, Prox-1, PDPN, and VEGFR3. NLVs frequently coil around blood vessels, and putative connections between vascular structures were observed. Notably, some NLVs traverse multiple cortical layers and display distinct orientation patterns that vary across cortical laminae.

This discovery reveals a previously “hidden” vascular network in the brain parenchyma and raises the possibility that an abundant, highly organized system of nanoscale tubular structures may provide an efficient conduit for metabolic waste clearance. Such a system could represent a critical, previously unrecognized component supporting the brain’s extraordinary metabolic demands.

## Introduction

Blood and lymphatic systems are two major circulatory systems in our body. The blood system runs through every organ, and most large organs, such as skin, liver, lung, heart, and kidney, also have a parallel lymphatic system, which is a one-way system transporting fluid from tissues back to the central circulation. It was once believed that some organs, however, are devoid of a lymphatic system, including the brain, eye, and spinal cord, due to immune privilege that prevents inflammation ^1–3^. However, this perspective has gradually been overturned due to several seminal breakthroughs ^4–10^. For example, in one groundbreaking study in 2012, Nedergaard and Iliff et al. showed that cerebrospinal fluid (CSF) flows into the brain parenchyma via perivascular spaces (PVS) around arteries, facilitated by aquaporin-4 (AQP-4) channels formed by astrocyte endfeet ^4^. This landmark study was the first to provide evidence for a lymphatic system within the brain. As the system relies on the function of glial cells, the lymphatic system in the brain was termed a “glymphatic” system. Later, in other landmark studies in 2015, Kipnis and Alitalo et al. revealed that meningeal lymphatic vessels (mLVs) exist in the dura mater surrounding the brain and can drain CSF and waste materials to peripheral lymph nodes ^5,11^, In 2017, the existence of meningeal lymphatics was confirmed in human autopsy samples ^12^. In 2019, Koh et al. demonstrated that meningeal lymphatics in the basal part of the skull are also crucial for the drainage of CSF ^7^. In 2020, Nedergaard et al. found that retina also contains glymphatic systems^13^.

Blood system consists of large vessels and small vessels, e.g. capillaries, and the artery blood exchanges with vein blood through capillary networks that are formed by abundant small vessels with diameters greater than 5 μm. Like the blood vasculature, the peripheral lymphatic system is composed of both large vessels and capillaries ^14–16^. In contrast, reports of capillary-level lymphatic vessels within the brain are exceedingly rare ^17,18^, and descriptions of dense lymphatics in the brain parenchyma are virtually absent. In 2023, Abumaria and Chang et al. discovered the presence of lymphatic vessels with diameters in the range of 5 -14 μm in deep brain areas such as the hippocampus and cortex; however, these structures were sparse ^18^. Therefore, it remains a major open question whether the brain, like peripheral organs, also possesses lymphatics that are formed by abundant capillary-like lymphatic vessels.

In this letter, we report the discovery of a totally new type of lymphatic vessel structure in the brain with diameters of 500-1000 nanometers with very high abundance, which forms webs of lymphatic-like system across the whole brain. We termed the new vessels “nano-scale lymphatic-like vessels (NLVs)”. These NLVs may function as structural conduits for neurofluid transport and waste clearance.

## Results

### Discovery of nano-scale abundant “hidden” vessels

While our initial goal was to obtain high-resolution images of Aβ plaques, we serendipitously discovered unique vessel-like structures in the cortical region, characterized by a hollow lumen and brightly labeled walls. This finding was made possible by the unique combination of expansion microscopy (ExM) technology and CRANAD-3, a highly fluorescent small-molecule probe (Fig.1A) ^19,20^. ExM not only provides high-resolution images without the need for super-resolution technologies such as STED or STORM, but also significantly increases the signal-to-noise ratio (SNR) by making the tissue nearly transparent and reducing scattering after the expansion process ^19,21,22^. CRANAD-3 is a highly responsive fluorescence probe for beta-amyloid (Aβ) that was designed by our group nearly a decade ago. The probe is sensitive to not only monomeric, oligomeric, but also fibrillar Aβ species^20,23,24^. As Aβ species are constantly cleared out from brain via glymphatic system^4,13,25–29^, the visualization of NLVs is likely attributable to CRANAD-3’s ability to label Aβ species deposited along glymphatic pathways.

**Figure. 1.**
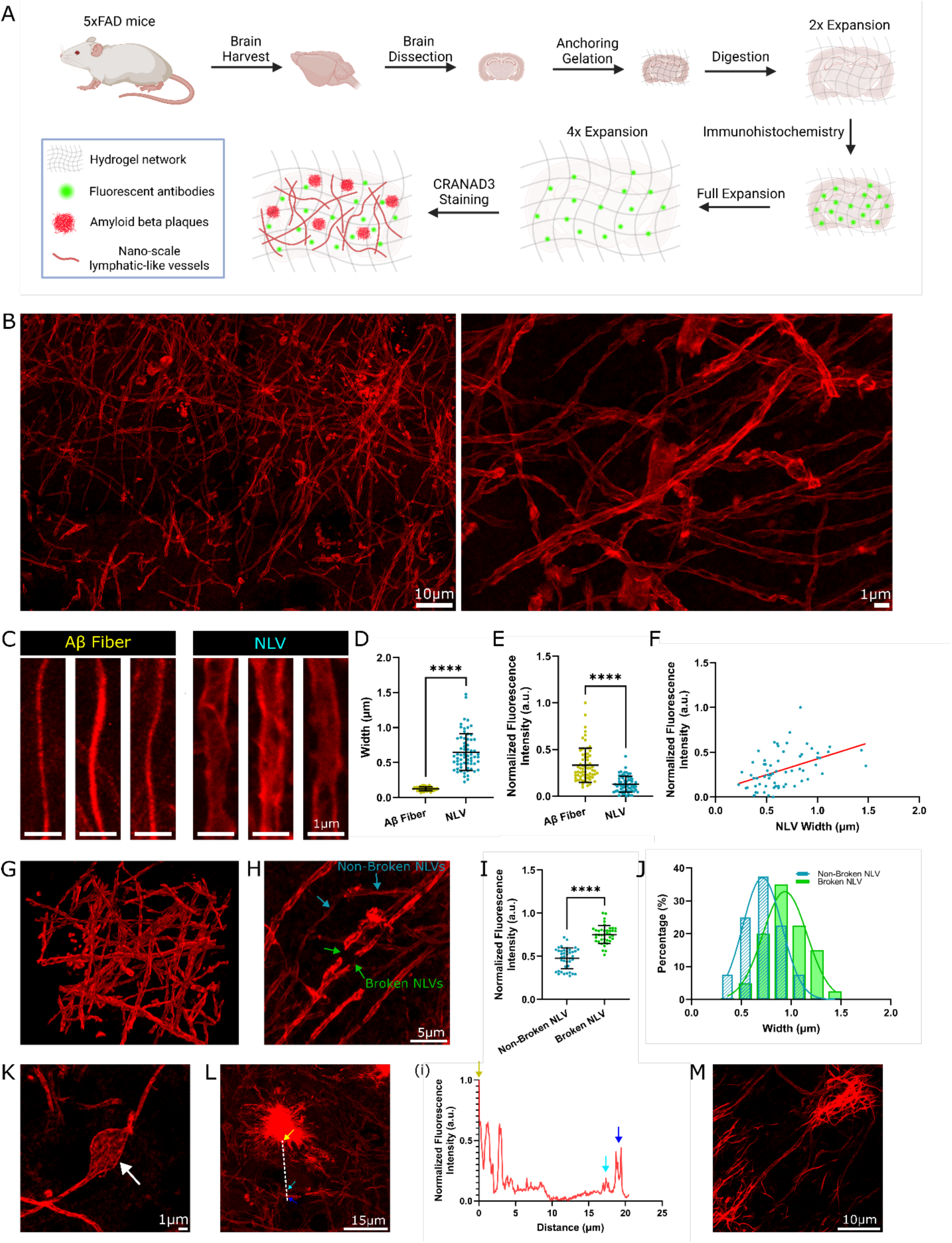
Discovery of nano-scale lymphatic-like vessels (NLVs) in brain parenchyma. (**A**) Experimental scheme of expansion microscopy imaging on mouse brain sections with CRANAD-3 and immunolabeling. **(B)** Representative expansion microscopy (ExM) images showing abundant NLVs (left panel) in the mouse brain parenchyma stained with CRANAD-3. The hollow feature can be clearly observed in the right panel. **(C)** ExM images showing three representative Aβ fibrils and NLVs. **(D)** Quantification of the widths of Aβ fibrils (n = 65) and NLVs (n = 65) (P < 0.0001). **(E)** Quantification of normalized fluorescence intensities of Aβ fibrils (n = 65) and NLVs (n = 65) (P < 0.0001). Fluorescence intensities were normalized by subtracting the minimum value of the two groups and then divided by the diSerence between the maximum and minimum values. a.u., arbitrary units. **(F)** Relationship between normalized fluorescence intensity and width of NLVs (n = 65, R^2^ = 0.2170, linear regression fitting). **(G)** Three-dimensional ExM reconstruction of NLVs. **(H)** ExM image showing non-broken (blue arrows) and broken (green arrows) NLVs. **(I)** Quantitative analysis of normalized fluorescence intensities of non-broken (n = 40) and broken (n = 40) NLVs (P < 0.0001), revealing higher fluorescence intensities of broken NLVs. **(J)** Frequency distribution of non-broken and broken NLV widths (n = 40 per group; R^2^ = 0.9976 for non-broken NLV group; R^2^ = 0.9568 for broken NLV group; Gaussian nonlinear regression), demonstrating larger diameters of broken NLVs. **(K)** ExM image showing burgeon bulbs on an NLV. White arrow points to a swollen part of the NLV. **(L)** NLVs surrounding an Aβ plaque from an ExM sample, demonstrating a void area around the plaque and squeezed NLVs. (i) Normalized fluorescence intensity profile of CRANAD-3 of the dashed line in (L); the yellow arrow indicates the Aβ plaque boundary, and the cyan and blue arrows mark two NLVs. The line plot shows an NLV-avoid area of about 15 μm length. **(M)** A radial plaque appearing at the end of broken NLVs. All images in Figure 1 were obtained from ExM-processed brain samples stained with CRANAD-3. For all scatter plots, data are presented as mean ± standard deviation (SD), and P values were calculated using nonparametric Mann-Whitney tests. ****P < 0.0001. Scale bars, 10 μm for left panel of (B), 1 μm for right panel of (B); 1 μm for (C); 5 μm for (H); 1 μm for (K); 15 μm for (L); 10 μm for (M).

Using 4×ExM combined with CRANAD-3 in AD mouse brain tissue, we observed unique vessel-like (tubular) structures in the cortical region, characterized by a hollow lumen with brightly labeled walls (Fig.1B,C and Movie 1 and Movie 2). These hollow vessels, with a diameter ranging from 500 to 1000 nm, are clearly distinct from Aβ fibrils (Fig.1D), which are abundant around the plaque core (SI Fig.1A, B) with observable diameter of 200-300 nm (Fig.1D, E). In addition, the observed NLVs’ fluorescence intensity is much lower than that of Aβ fibrils (Fig.1E). Notably, distant from plaques, we observed abundant vessels with similar features, forming web-like structures with both parallel and intersecting (including perpendicular) orientations (Fig. 1B,G). In addition, such unique vessels were observed in multiple brain areas and different mouse brain tissues (SI Fig.1C).

Several other notable features are also clearly visible in the images obtained with CRANAD-3. First, 3D imaging showed that most of the nano-vessels are independent of each other, and the nano-vessels do not have apparent tree-like branches, even when the vessels cross each other. (Fig.1G, and SI Fig.1). Second, the bright vessels have punctate bright patches along the vessels (Fig.1H). Third, notably, we found that compared to the dim NLVs, the brightest vessels seem more prone to breakage (Fig.1H, I, J). Fourth, we observed burgeon bulbs along the vessels, and bright patches could be observed in the burgeon bulbs (Fig.1K). Moreover, a notable feature is the presence of void zones lacking nano-vessels surrounding large plaques (Fig.1L). Lastly, we found some radial plaques at the end of broken nano-vessels, which may indicate that broken nano-vessels may facilitate plaque formation (Fig.1M).

The unique diameter size range (500-1000 nm) excludes most of the vessels and vessel-like structures, such as large blood vessels and canonical lymphatic vessels ^30^. It also excludes blood capillaries, as their smallest sizes are about 4-6 microns to allow blood cells to flow through ^31^. Additionally, these structures do not correspond to previously reported lymphatic capillaries, which have minimum diameters exceeding 5 microns ^18^. Although pain fibers (unmyelinated axon) have diameters of 0.1–1 μm, they are not abundantly present in the brain ^32^. After a systematic survey of vessel-like structures with such size features, we speculated that the nano-scale vessels are “hidden” lymphatics.

### Validation of lymphatic-like features

To validate that the nano-vessels are lymphatics, we first used LYVE-1, the most commonly used marker for lymphatics ^33^, with its antibody both without and with ExM. We observed some features that are close to sub-micron diameter size of fragmented lines that are stretched from the surface of the cortex to inner layers. However, these lines are made of punctate LYVE-1 antibody staining, and they do not clearly show the hollow features of vessels (SI Fig.2A). This punctate appearance is consistent with reported references ^18,34^; however, intriguingly, no references mentioned such linear features of sub-micron size. In addition, it is obvious that there are abundant random puncta in the areas without linear features, and it seems that the puncta are not merely noise (SI Fig.2A). To investigate whether ExM can provide better resolutions for the expected vessel features, we used 4x ExM with LYVE-1 antibody staining. After ExM, we totally lost the nano-scale vessel feature (SI Fig. 2B). It appears that the punctate LYVE-1 staining pattern was insufficient to reveal the “hidden” nano-vessel structures in our study. This non-continuous (punctate) labeling may also explain why other groups failed to identify these nano-scale vessels, given that LYVE-1 is widely regarded as the gold-standard marker for lymphatic characterization.

**Figure. 2.**
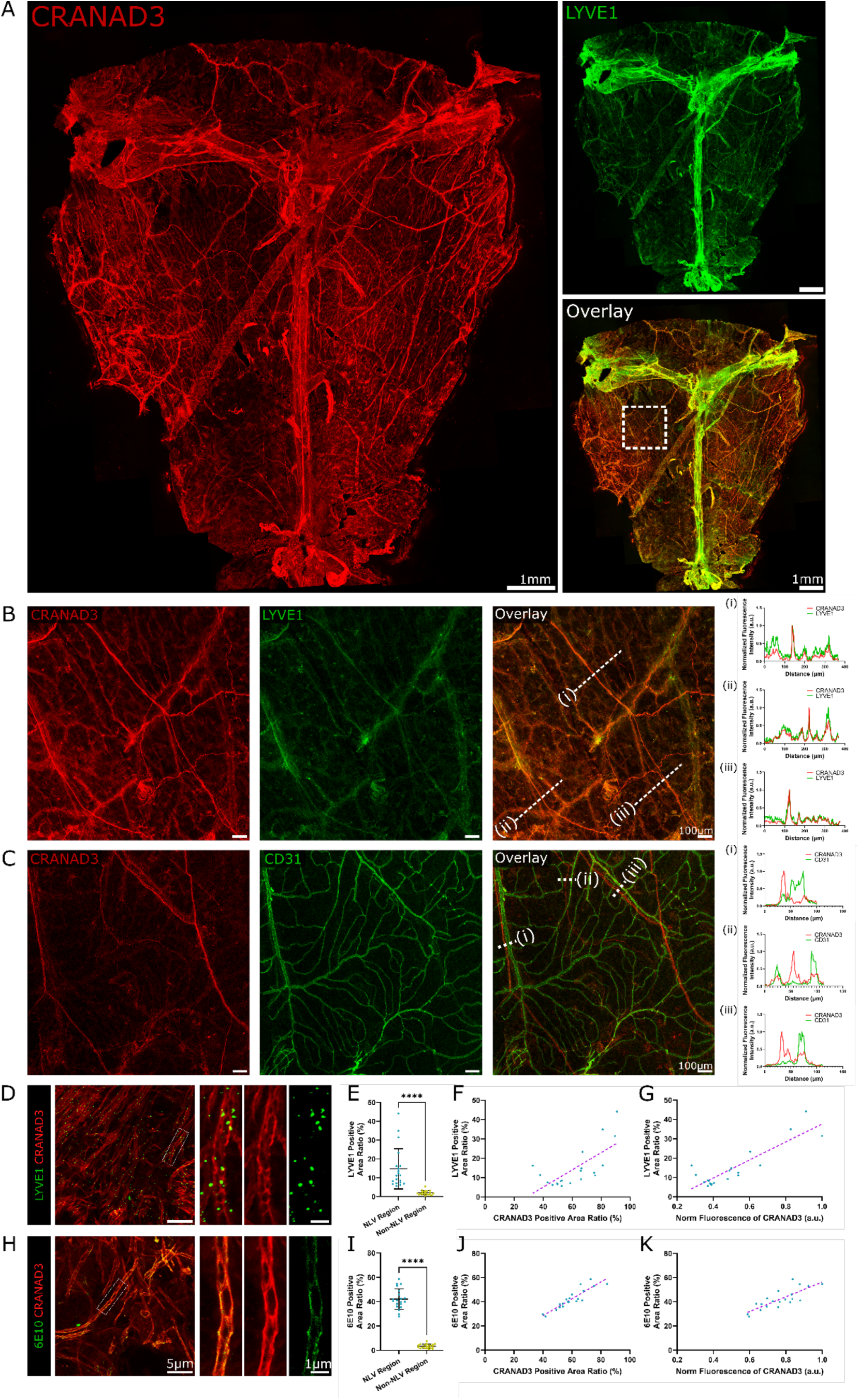
Validation of lymphatic-like features of NLVs in meninges and parenchyma of 5xFAD mouse brain. **(A)** Non-ExM images of whole-mounted WT mouse meninges stained with CRANAD-3 (red) and lymphatic-specific marker LYVE-1 (green), showing strong co-localization. **(B)** Magnified view of the boxed region in the overlay panel of (A) showing co-localization of CRANAD-3 and LYVE-1 signals. (B) (i-iii) Normalized fluorescence intensity profiles of LYVE-1 and CRANAD-3 of the dashed lines in the overlay panel of (B). **(C)** Non-ExM images of partial whole-mounted 5xFAD AD mouse meninges stained with CRANAD-3 (red) and blood vessel-specific marker CD31 (green). (C) (i-iii) Normalized fluorescence intensity profiles of CD31 and CRANAD-3 of the dashed lines in the overlay panel of (C), confirming no apparent overlap between CRANAD-3 and CD31 staining in meninges. **(D)** Representative ExM image of 5xFAD mouse brain section co-stained with CRANAD-3 and LYVE-1. Right panels show magnified views of the boxed region in the left panel. **(E)** Quantification of LYVE-1 normalized fluorescence intensities at NLV areas (n = 20) and non-NLV areas (n = 20) (P < 0.0001). **(F)** Relationship between LYVE-1-positive area ratio and its correlation with CRANAD-3-positive areas at NLV regions (n = 20) (R^2^ = 0.4932, linear regression fitting). To quantify positive area ratio, a threshold was set; any pixel with a value above it was classified as positive. **(G)** Relationship between LYVE-1-positive area ratio and normalized fluorescence intensity of CRANAD-3 at NLV regions (n = 20) (R^2^ = 0.7686, linear regression fitting). **(H)** ExM image of 5xFAD mouse brain section co-stained with CRANAD-3 and 6E10. Right panels show magnified views of the boxed region in the left panel. Patchy 6E10 signals align very well with the vessel highlighted by CRANAD-3. **(I)** Quantification of 6E10 normalized fluorescence intensities at NLV areas (n = 20) and non-NLV areas (n = 20) (P < 0.0001). **(J)** Relationship between 6E10-positive area ratio and its correlation with CRANAD-3-positive areas at NLV regions (n = 20) (R^2^ = 0.8194, linear regression fitting). **(K)** Relationship between 6E10-positive area ratio and normalized fluorescence intensity of CRANAD-3 at NLV regions (n = 20) (R^2^ = 0.6472, linear regression fitting). For all scatter plots, data are presented as mean ± standard deviation (SD), and P values were calculated using nonparametric Mann-Whitney tests. ****P < 0.0001. Scale bars, 1 mm for (A); 100 μm for (B) and (C); 5 μm for left panels of (D) and (H), 1 μm for right panels of (D) and (H).

Since the discovery of glymphatic system in brains in 2012 and the report of lymphatics in meninges in 2015, the lymphatic system in meninges has been firmly validated ^2,5,7,35–38^. The lymphatic vessels are easily identified through various imaging methods without tissue expansion or super-resolution imaging. Among the imaging methods, LYVE-1 antibody staining is still the gold standard. In this regard, we reasoned that if the staining patterns of CRANAD-3 and LYVE-1 are similar, this would strongly suggest that CRANAD-3 is capable of visualizing lymphatic vessels both in meninges and parenchyma. With these considerations, we co-stained mouse meninges with LYVE-1 and CRANAD-3 without tissue expansion (hereafter referred to as “non-ExM.”). Our data demonstrate that CRANAD-3 provided similar staining patterns to LYVE-1 with the whole mounted meninges (Fig.2A, B). Both CRANAD-3 and LYVE-1 showed strong signals from the trunk regions of superior sagittal sinus (SSS) and transverse sinus (TS), and both revealed details of vessels in the off-trunk regions. As expected, the signals showed excellent co-localization (Fig.2B). Although the overall staining patterns are similar, there are several notable differences. As widely reported, LYVE-1 showed much stronger signals from the trunks, while the signals in the off-trunk regions are relatively weak (Fig.2A)^2,5,7,35,36^. Notably, we noticed that CRANAD-3 provided strong staining for both trunk and off-trunk regions (Fig.2A, B).

As meninges also contain abundant blood vessels, it is important to confirm that the CRANAD-3 signals are not from blood vessels. To this end, we co-stained the whole mounted meninges with CRANAD-3 and CD31, a widely used marker for blood vessels ^39,40^. As expected, the staining patterns from CRANAD-3 and CD31 are distinct from each other. There are no significant overlaps between them in either trunk or off-trunk regions (Fig.2C and SI Fig.2C). Taken together, the above results strongly indicated that CRANAD-3 can stain lymphatic vessels but not blood vessels.

Encouraged by CRANAD-3’s capacity for staining lymphatics in meninges, we re-examined the images of the expanded cortex tissue that were co-stained with LYVE-1 and CRANAD-3. Notably, after careful examination, we observed that LYVE-1 positive puncta align along the NLVs highlighted by CRANAD-3 (Fig. 2D). To further evaluate the association between CRANAD-3 and LYVE-1, we first quantified the positive areas of LYVE-1 that aligned with vessels and intensities from areas that had no clear vessel association (non-NLV areas). We found that the averaged LYVE-1 signals were about 7.5-fold higher in the NLV-associated areas than in the non-NLV areas (Fig.2E). To further investigate the association, we performed a correlation between CRANAD-3 positive areas and LYVE-1 positive areas for NLVs and found that the correlation could reach R^2^ = 0.5, indicating that CRANAD-3 and LYVE-1 are correlated (Fig.2F). In addition, we found an excellent correlation (R^2^ = 0.8) between CRANAD-3 intensities and LYVE-1 positive area ratios (Fig.2G). Taken together, although LYVE-1 could not highlight the contours of NLVs, it showed strong associations with NLVs that were outlined by CRANAD-3.

Prox-1 is a homeodomain transcription factor critical for the development and maintenance of lymphatic endothelial cells (LECs), and is another commonly used lymphatic marker ^41,42^. In this regard, we used its antibody to investigate whether Prox-1 can be used to visualize the hidden nano-vessels. Again, although numerous studies have used different Prox-1 antibodies to image the glymphatic system ^41,42^, none of the studies uncovered the unique nano-tubular structures observed here. In some cases, advanced optical imaging, such as light sheet with a combination of tissue clearance, which also increases resolution by reducing scattering, was used, but it only revealed lymphatics with diameters > 5 microns ^18^. We then asked whether the combination of ExM/CRANAD-3/Prox-1 could facilitate the validation of nano-lymphatics. Again, ExM/Prox-1 was unable to reveal clear contours of NLVs. However, we observed that Prox-1 positive puncta align along with the nano-vessels highlighted by CRANAD-3 (SI Fig 2D). We further performed correlation studies that are similar to LYVE-1 and found that Prox-1 was highly associated with NLVs (SI Fig.2E-G).

Podoplanin (PDPN) is a transmembrane glycoprotein highly expressed in LECs, and PDPN has often been used as a marker for lymphatic imaging ^43,44^. Similar to LYVE-1 and Prox-1 staining with ExM, PDPN also showed punctate staining. Similar to other lymphatic markers, the PDPN puncta align with NLVs (SI Fig.2H-K). In addition, VEGFR3 is another widely used marker for characterizing lymphatics ^45,46^. We performed similar ExM staining with VEGFR3/CRANAD-3 and found that the VEGFR3 signals also align with the NLVs (SI Fig.2L-O).

It is well documented that Aβs are transported out of the parenchyma via glymphatics ^4,13,25–29^. Hence, it is reasonable to speculate that portions of Aβ may deposit on the wall of the vessels, due to the sticky property of Aβs. In this regard, we stained the expanded cortex tissue with two Aβ antibodies: 6E10, a pan-Aβ antibody ^47^, and 12F4, an Aβ42 specific antibody ^48^. We found that 6E10 staining co-localized strongly with CRANAD-3 signals and could delineate NLV contours; however, its staining quality was inferior to that of CRANAD-3 (Fig.2H). We also performed correlation studies with signal intensity, 6E10 positive area ratios, and found that 6E10 staining was tightly correlated to CRANAD-3 (Fig.2I-K). Notably, in contrast to the staining with LYVE-1 and PDPN, and Prox-1, the antibody staining can highlight the vessel feature, instead of punctate features. The continuous staining feature of antibody is similar to CRANAD-3, suggesting that the capacity of CRANAD-3 staining is probably due to its known binding to Aβ deposits on NLVs. Because CRANAD-3 can label Aβ monomers, dimers, oligomers, and aggregates, whereas 6E10 primarily recognizes aggregated Aβ, this may explain CRANAD-3’s superior labeling of NLV walls. In addition, we performed the same experiments with 12F4 antibody, and we found that its signals also align with the CRANAD-3 positive NLVs as puncta or short stretches, but its continuity was inferior to that with 6E10 staining (SI Fig.2P-S). Intriguingly, these punctate stretches of 6E10 were described as “nano clusters” in 2022 ^22^. With a careful survey, we found that a linear pattern could be observed from the reported nanoclusters. Taken together, CRANAD-3’s labeling of NLVs is consistent with its established ability to stain Aβ in its various forms.

### ExM with WT mouse brain and human AD brain slides

To investigate whether such nano-vessels can be found in wild-type (WT) mouse brains, we used the same expansion protocol and CRANAD-3 for staining. As expected, we found that there were also abundant nano-vessels but with apparently weaker CRANAD-3 signals, compared to AD mouse brain (Fig.3A). The ability of CRANAD-3 to visualize NLVs is likely attributable to its high sensitivity to rat and mouse Aβ species, as we reported previously in 2015 ^20^. Although rodent Aβ has a much lower propensity than human Aβ to accumulate along nano-vessels, it can nevertheless do so because of its highly hydrophobic and adhesive nature. Thus, the observed images most likely reflect CRANAD-3 staining of rodent Aβ, consistent with our prior demonstration that CRANAD-3 is responsive to rat Aβ ^20^. As expected, using the same ExM imaging protocol, we also observed abundant NLVs in the cortex of brain slides of an AD patient (Fig.3A, right panel). These results strongly suggest that the existence of NLVs is a generic phenomenon across different species. Interestingly, we found there were less broken NLVs in WT brain, compared to AD mouse brain and human AD brain (Fig.3B, D, F), while all broken NLVs showed higher CRANAD-3 intensity, compared to non-broken NLVs (Fig.3C, E, G). We then compared the fluorescence intensity and SNRs and found that the WT brain showed the lowest SNRs and the human AD subject brain had the highest SNR, while they showed a similar trend with fluorescence intensity (Fig.3H, J, K). Regarding the diameters, the human AD brain showed the largest sizes while slightly larger sizes were obtained in AD mouse brain than in WT brains (Fig.3L). We also analyzed the percentage of the CRANAD-3 positive area under the same imaging conditions and found that the human AD brain had the highest percentage (Fig.3I, N). Notably, we observed the more frequent appearance of burgeons on the NLVs in human AD brain (Fig.3O, P). In addition, we noticed that several NLVs bundled together in the human AD brain (Fig.3Q).

**Figure. 3.**
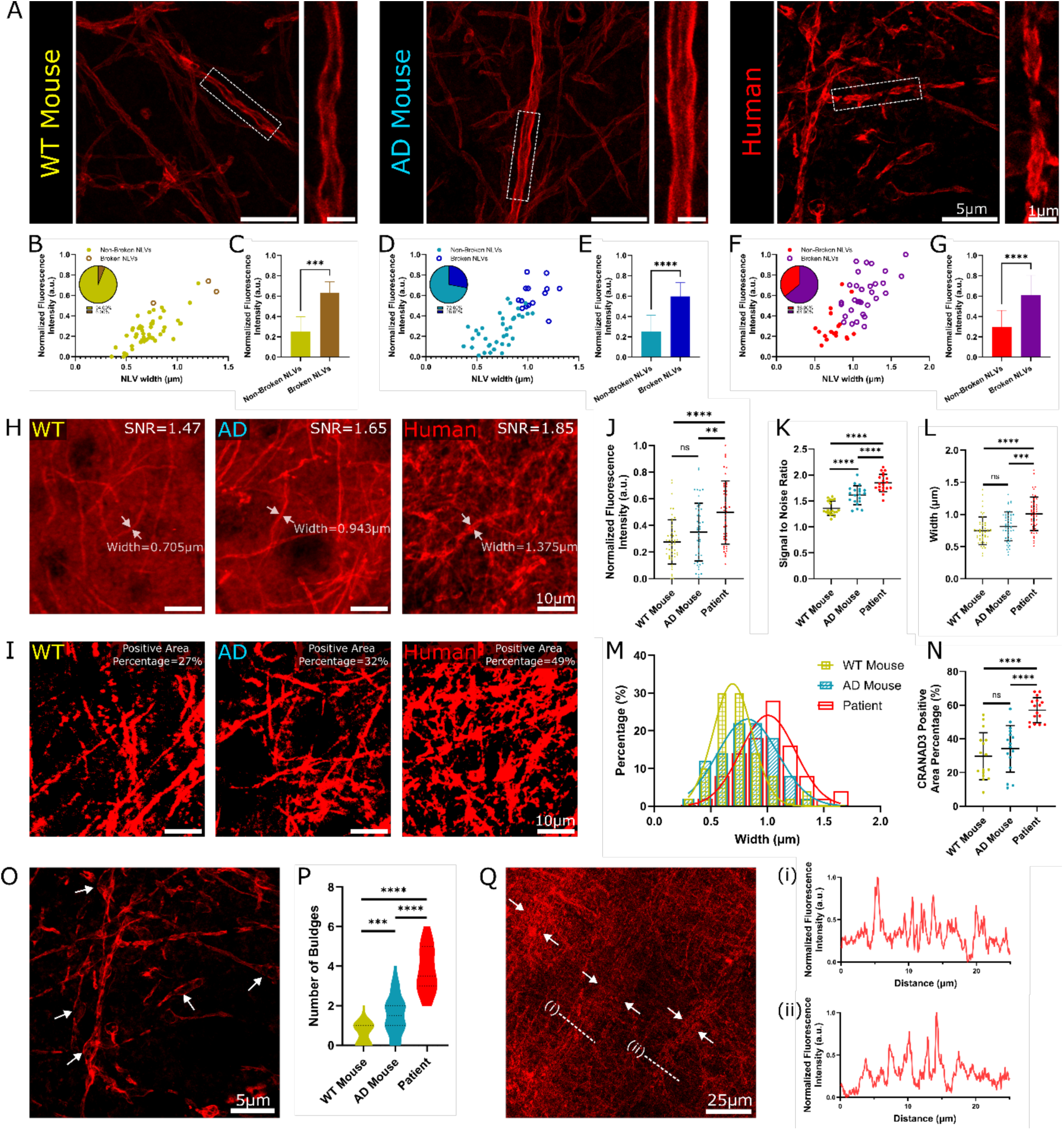
Comparison of NLV features in WT mice, 5xFAD mice, and patient brain. **(A)** Representative ExM images of WT mice, 5xFAD mice, and patient brain sections stained with CRANAD-3. Magnified images of the boxed regions in each panel are shown to the right of each panel. **(B, D, F)** Relationship between normalized fluorescence intensity of CRANAD-3 and NLV width in WT mice group (B), AD mice group (D), and patient group (F) (n = 50 per group). Pie charts show the percentage of broken NLVs among all NLVs in each group: WT (6%), AD (28%), and patient (64%). **(C, E, G)** Quantification of normalized CRANAD-3 fluorescence intensities at broken-NLVs and non-broken NLVs in WT mice group (C), AD mice group (E), and patient group (**G**). Broken NLVs showed significantly higher fluorescence [P = 0.0007 for (C); P < 0.0001 for (E) and (G)]. (C), (E), and (G) use the same dataset with (B), (D), and (F), respectively. **(H)** Representative ExM images stained with CRANAD-3 from the three groups, with typical signal-to-noise ratios (SNRs) and NLV widths indicated. **(I)** ExM CRANAD-3-stained samples, showing typical CRANAD-3-positive area ratios. **(J)** Quantification of normalized fluorescence intensities of CRANAD-3 in 3 groups (n = 50 per group; P = 0.0022 for AD mouse vs. patient comparison; P < 0.0001 for WT mouse vs. patient comparison). **(K)** Quantification of SNRs of CRANAD-3 across the three groups (n = 50 per group; P < 0.0001 for all three comparisons). **(L)** Quantification of NLV widths across the three groups (n = 50 per group; P = 0.0002 for AD mouse vs. patient; P < 0.0001 for WT mouse vs. patient comparisons). **(M)** Percentage histogram of NLV widths across three groups (n = 50 per group; Gaussian nonlinear regression, R^2^ = 0.9598 for WT mice group; R^2^ = 0.9652 for AD mice group; R^2^ = 0.9256 for patient group). **(N)** Quantification of CRANAD-3 positive area percentages across the three groups (n = 50 per group; P < 0.0001 for AD mouse vs. patient; P < 0.0001 for WT mouse vs. patient comparisons). **(O)** ExM CRANAD-3-stained image from patient brain slice. White arrows show swollen NLVs. **(P)** Quantification of the number of burgeon bulbs in the images of the three groups (n = 30 regions; P = 0.0001 for WT mouse vs. AD mouse; P < 0.0001 for AD mouse vs. patient; P < 0.0001 for WT mouse vs. patient comparisons). **(Q)** ExM image of patient brain slice, showing NLV bundles extending in a straight and parallel manner. (i) and (ii) Normalized fluorescence intensity profiles of CRANAD-3 of the dashed lines in (Q). For all scatter plots, data are presented as mean ± standard deviation (SD), and P values were calculated using nonparametric Mann-Whitney tests. n.s., not significant; **P < 0.01; ***P < 0.001; ****P < 0.0001. Scale bars, 5 μm for left panels of (A), 1 μm for right panels of (A); 10 μm for (H) and (I); 5 μm for (O); 25 μm for (Q).

After demonstrating that WT-NLVs can also be visualized with CRANAD-3, we next examined WT brain sections to assess whether the punctate staining of LYVE-1, Prox-1, VEGFR3, and PDPN aligns with the nano-vessels highlighted by CRANAD-3/ExM. Indeed, the staining patterns of these markers in WT brains were similar to those observed in AD mouse brains (SI Fig.3A-D).

### Exclusion of blood vessels, neuron axons, dendrites, and collagens

To exclude the possibility that the NLVs represent other linear fibrils (such as collagen fibrils and neuronal axons) and tubular structures/vessels (including microtubules in neurons and blood capillaries), we performed ExM with several immuno-histological markers. To exclude the possibility of axons, we used an antibody against Ankyrin-G (AnkG), a key protein in the axon initial segment (AIS) of a neuron ^49,50^, to stain the expanded tissue, while CRANAD-3 was used to visualize the NLVs. As expected, there were no clear outlines of vessels by AnkG. Although we observed punctate spots, no strong correlation could be established (SI Fig.4A). MAP2, microtubule-associated protein 2, is a neuron-specific protein found primarily in the soma and dendrites^51^. With MAP2 antibody staining, we could rule out that microtubules overlap with NLVs labeled with CRANAD-3 (SI Fig.4B). As NLVs could appear as a fibrillar feature in low-magnification microscopic images, and such fibrillar feature could be assembled into fibrils in extracellular matrix (ECM). In this context, we examined collagen I, a major component of the brain extracellular matrix (ECM) ^52^, which exhibits a fibrillar staining pattern, to assess whether its distribution resembles that of CRANAD-3. As expected, collagen I and CRANAD-3 showed distinct and nonoverlapping staining patterns in expanded cortical tissues (SI Fig.4C).

**Figure. 4.**
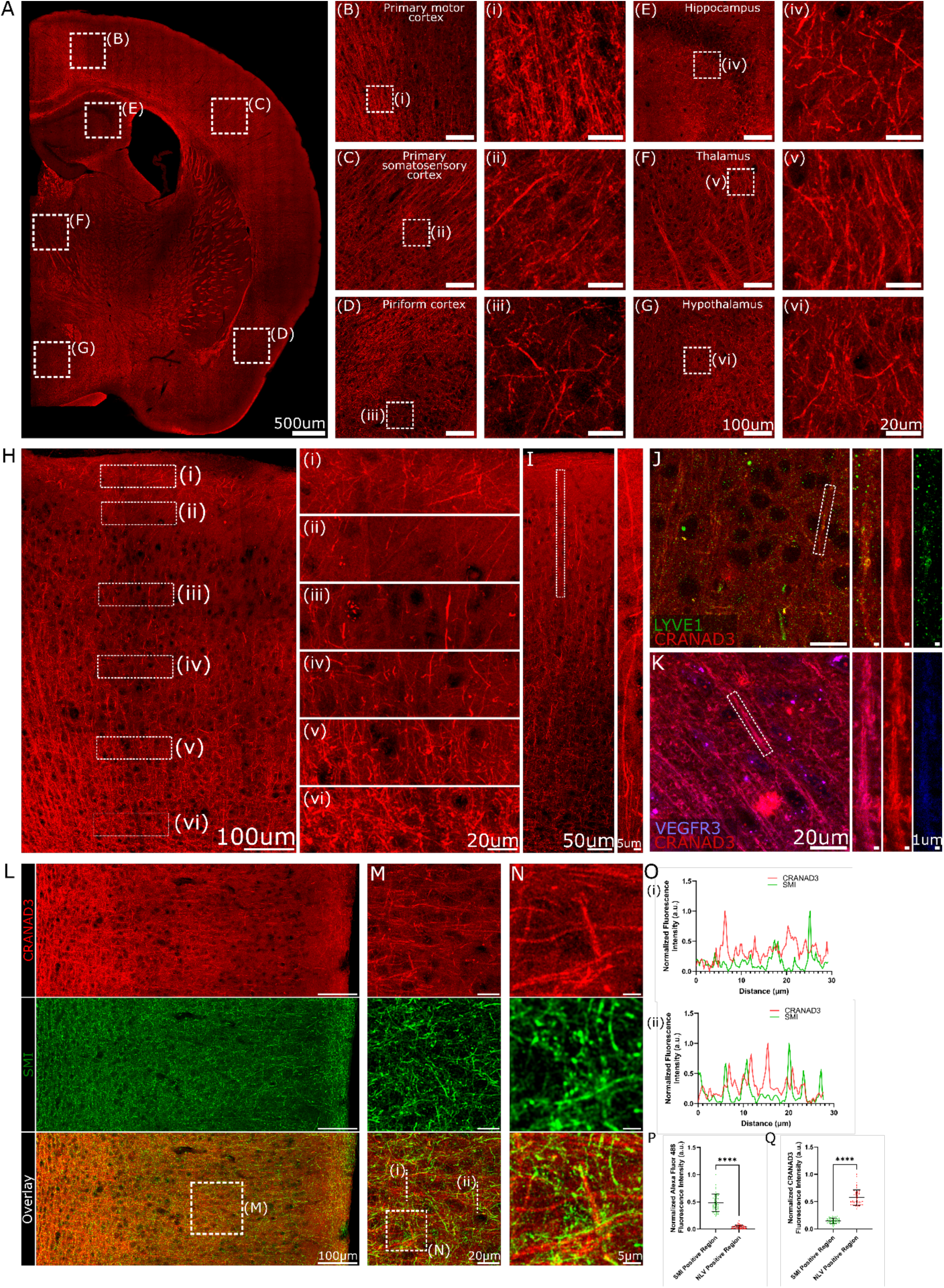
Non-ExM imaging of NLVs with CRANAD-3. **(A)** non-ExM image of half of a WT mouse brain section acquired using a water-compatible objective in PBS. **(B-G)** Magnified views of different brain regions indicated in (A), including primary motor cortex (B), primary somatosensory cortex (C), piriform cortex (D), hippocampus (E), thalamus (F), and hypothalamus (G). Higher magnification views o**f** the boxed regions are shown in (i-vi), respectively. **(H)** High-resolution image of primary somatosensory cortex and zoomed-in views of diSerent layers (i – vi). **(I)** Representative image showing an individual NLV that extends across diSerent layers. Right panel shows the magnified view of the boxed region in the left panel. **(J-K)** Non-ExM co-immunostaining of LYVE-1 (J) and VEGFR3 (K) with CRANAD-3. LYVE-1 puncta align with NLV structure, and VEGFR3 shows a faint contour of NLVs. Magnified views of the boxed regions are shown in the right panels. **(L-N)** Non-ExM sample co-immunostaining of SMI-31 with CRANAD-3 in the cortex of WT mouse brain slices. (M-N) Magnified images of the boxed regions in (L) and (M), respectively. **(O)** Normalized fluorescence intensity profiles of SMI-31 and CRANAD-3 of the dashed lines shown in (M). **(P-Q)** Quantitative analysis of normalized SMI-31 and CRANAD-3 fluorescence intensities in SMI-31-positive areas and NLV-positive (CRANAD-3) areas (n = 50 per group; P < 0.0001 for both analyses). The distinct staining patterns confirmed that the CRANAD-3 signal does not originate from neuron axons. For all scatter plots, data are presented as mean ± standard deviation (SD), and P values were calculated using nonparametric Mann-Whitney tests. ****P < 0.0001. Scale bars, 500 μm for (A); 100 μm for (B-G), 20 μm for (i-vi); 100 μm for (H), 20 μm for (i-vi); 50 μm for left panel of (I), 5 μm for right panel of (I); 20 μm for left panels of (J) and (K), 1 μm for right panels of (J) and (K); 100 μm for (L); 20 μm for (M); 5 μm for (N).

### NLV imaging with non-expanded brain tissues

Although we attempted to image NLVs in non-expanded brain tissue, the tissue-drying process prevented clear visualization of NLV contours with CRANAD-3. In contrast, when imaging in water or PBS buffer using a water-immersion objective at high magnification (40×–60×), NLVs were readily observable, albeit without a clearly defined hollow lumen (SI Fig.4D,E). The failure with dried tissue is likely due to high background signals of CRANAD-3, whose fluorescence intensity can be “turned-on” when it interacts to hydrophobic environments in dry tissues via non-specific binding. Using the non-ExM preparations, we again employed antibodies against AnkG and MAP2 to exclude the possibility that the signal originated from neuronal axons or dendrites. The distinct localization patterns observed confirmed that the CRANAD-3 signal does not arise from neuronal structures such as axons or dendrites (SI Fig.4D). We further confirmed that NLVs are distinct from small blood vessels, as CD31 and CRANAD-3 staining exhibited clearly different patterns (SI Fig.4E-J). Notably, while NLVs could be visualized in non-expanded wet tissue, the background signal was substantially higher compared to ExM. Consequently, the apparent abundance of NLVs in non-ExM image was much lower than that observed with ExM (SI Fig.5A-D). Additionally, we observed that commonly used mounting media for slide sealing can significantly interfere with CRANAD-3 staining. As an organic small molecule, CRANAD-3 can be dissolved or absorbed by the mounting media, causing image blurring—especially nano-scale structures.

**Figure. 5.**
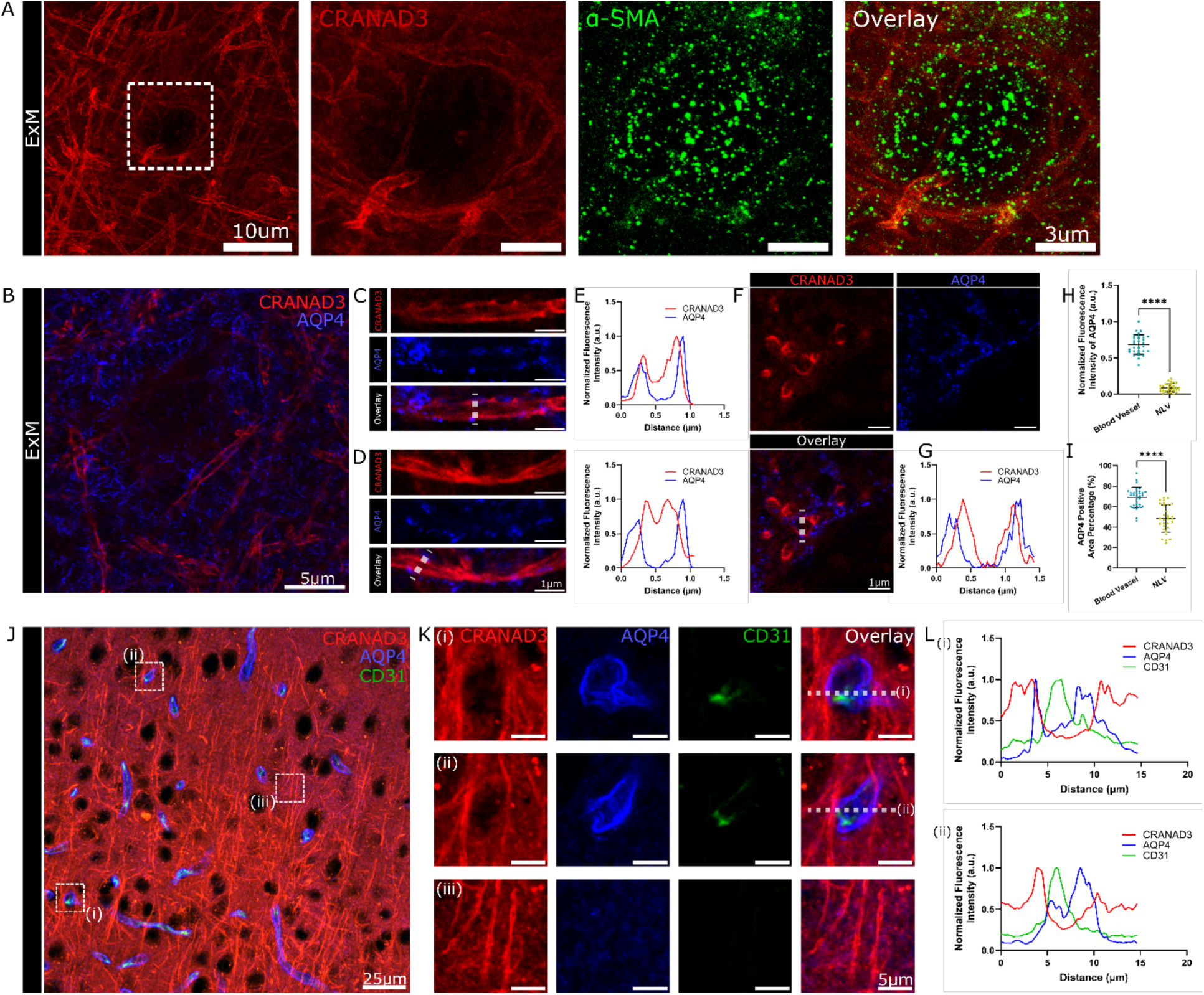
NLVs interact with blood vessels via aquaporin-4 channels. **(A)** ExM image of a WT mouse brain section co-stained with CRANAD-3 and α-SMA, demonstrating NLVs coiling around blood vessels marked by α-SMA positive pericytes. Magnified views of the boxed region are shown to the right. **(B)** ExM imaging of CRANAD-3 and aquaporin-4 (AQP-4) showing that NLVs intertwine with AQP-4. **(C-D)** Representative ExM images of individual NLVs, showing AQP-4 signals distributed on or adjacent to the NLVs. **(E)** Normalized fluorescence intensity profiles of CRANAD-3 and AQP-4 along the dashed lines in (C) and (D). **(F)** Cross-sectional view of ExM images of CRANAD-3 and AQP-4 signals. NLVs appear as holes here, and they are surrounded by AQP-4 signals. **(G)** Normalized CRANAD-3 and AQP-4 fluorescence intensity profiles of the dashed lines in the overlay panel of (F). **(H-I)** Quantitative comparison of normalized AQP-4 fluorescence intensities (H) and AQP-4-positive area percentage (I) around blood vessels and NLVs (n = 30 per group; P < 0.0001 for both analyses). Both NLVs and blood vessels are associated with AQP-4. **(J)** Non-ExM images of mouse cortex co-stained with CRANAD-3, AQP-4, and CD31. **(K)** Magnified views of the boxed regions in (J). **(L)** Normalized fluorescence intensity profiles of CRANAD-3, AQP-4, and CD31 along the dashed lines in (K). NLVs coiled around CD31-positive blood vessels with surrounding AQP-4-defined perivascular spaces. For all scatter plots, data are presented as mean ± standard deviation (SD), and P values were calculated using nonparametric Mann-Whitney tests. ****P < 0.0001. Scale bars, 10 μm for left panel of (A), 3 μm for right panel of (A); 5 μm for (B); 1 μm for (C) and (D); 1 μm for (F); 25 μm for (J); 5 μm for (K).

### Non-ExM and NLVs in different brain areas and cortical layers

We first performed non-ExM imaging on half of a brain slide and successfully visualized NLVs. Notably, their distribution patterns varied across brain regions (Fig. 4A–G). In the motor cortex, NLVs were bundled to form columnar structures (Fig. 4A, B), whereas in the primary somatosensory cortex, most NLVs were non-bundled (Fig. 4C). Interestingly, in layers 3 and 4 of the piriform cortex, NLVs appeared randomly arranged without a clear orientation (Fig. 4D). Notably, the NLV density in layer 1 of the piriform cortex was higher than in layer 1 of other cortical areas, with an orientation of approximately 45° relative to the cortical surface. NLVs were also identifiable in the hippocampus, thalamus, and hypothalamus (Fig. 4E–G). However, we noticed that CRANAD-3 could provide clear signals with fiber bundles of white matter (Fig.4A and SI Fig.1C), which is likely due to non-specific binding with white matter. Next, we performed higher-resolution non-ExM imaging of the primary somatosensory cortex. Remarkably, distinct NLV orientations were observed across different cortical layers (Fig.4H). In layer 1, most NLVs ran parallel to the cortical surface with moderately high density, though the SNR was low. In contrast, layer 2 exhibited low NLV abundance and low SNR, with only a few vessels detectable. Layers 3–4 showed NLVs oriented perpendicular to the cortical surface, with high abundance and relatively strong SNR; layer 3 was more densely populated than layer 4. In layers 5–6, NLVs formed a web-like net with both parallel and perpendicular orientations, maintaining high density. Interestingly, several NLVs were observed spanning from layer 1 through layer 3, suggesting long-range connectivity within the cortical NLV network (Fig.4I).

### Non-ExM with LYVE-1, Prox-1, PDPN and VEGFR3 staining

With the non-expanded brain slides, we attempted again to use the classical markers, such as LYVE-1, Prox-1, PDPN and VEGFR3, to image NLVs. As expected, all the immunostaining with antibodies against these markers failed to reveal clear NLV contours. However, except for PDPN, we found that LYVE-1, Prox-1, and VEGFR3 antibody could provide faint contours of NLVs (Fig.4J-K and SI Fig.4K, L, M). Given that LYVE-1, Prox-1, and VEGFR3 are essential markers for lymphatics, these results again suggest that NLVs are related to the lymphatic system.

### Non-ExM axon imaging with SMI-31 and NLV imaging with CRANAD-3

With non-ExM, we observed different patterns of NLV arrangements in different brain areas. Notably, it seems that the NLV arrangement patterns are similar to the arrangements of axons of neurons. To exclude the possibility that NLVs are axons, we performed staining with CRANAD-3 and SMI-31 antibody, the gold standard for assessing axonal integrity ^53^. As expected, the NLVs highlighted by CRANAD-3 are not axons, as the detailed staining patterns are distinct from each other (Fig.4L,M,N), and line plots and quantitative analysis also confirmed that CRANAD-3 signals are distinct from SMI-31 signals (Fig.4O,P,Q).

### NLV imaging with ExM and non-ExM with spinal cord

Similar to the brain, the spinal cord has traditionally been considered an immune-privileged site lacking lymphatics ^54^. However, recent studies have identified lymphatics in the spinal meninges ^34,55^. Whether NLVs exist within the parenchyma of the spinal cord remains unclear. To address this, we first applied ExM/CRANAD-3 and observed abundant NLVs in the spinal cord (SI Fig.6A). Non-ExM imaging similarly revealed easily identifiable NLVs, paralleling observations in the brain parenchyma (SI Fig.6B). In contrast, when performing comparable imaging on peripheral lymph nodes, such as deep cervical lymph nodes (dcLN), no NLVs were detected (SI Fig.6C, D).

**Figure. 6.**
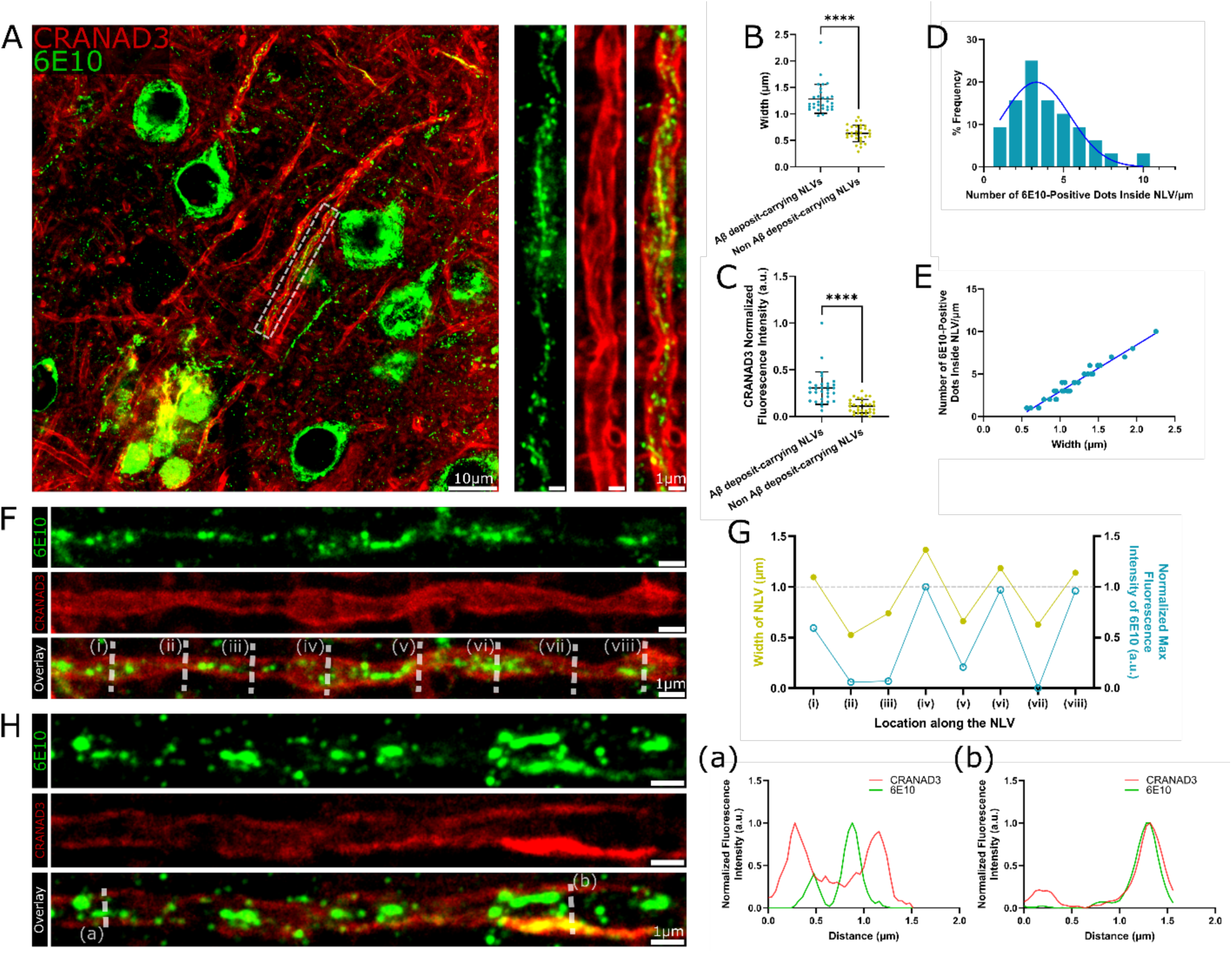
Aβ transportation and deposition within NLVs. (**A**) Representative non-ExM image of AD (5xFAD) mouse brain section stained with CRANAD-3 and 6E10. Magnified views of the boxed regions show 6E10-positive Aβ deposits located within NLVs. **(B)** Comparison of NLV diameter with and without Aβ deposits inside, demonstrating that NLVs containing Aβ deposits exhibit significantly larger widths (n = 30; P < 0.0001). **(C)** Quantification of normalized CRANAD-3 fluorescence intensities of NLVs with and without Aβ deposits inside (n = 30; P < 0.0001). **(D)** Frequency distribution of 6E10-positive Aβ puncta per micrometer of NLV length (n = 32; R^2^ = 0.8792; Gaussian nonlinear regression). **(E)** Correlation analysis between NLV width and numbers of Aβ deposit puncta inside NLVs per micrometer (n = 32; R^2^ = 0.9555; linear regression fitting). **(F)** Image of single NLV containing 6E10-positive Aβ deposits. **(G)** Quantification of NLV diameters and normalized maximum fluorescence intensities of 6E10 signals at eight diSerent locations (i-viii) along the NLV in (F). Regions with stronger 6E10 signals exhibit increased NLV thickness (>1 μm), whereas regions lacking or showing low 6E10 signals remain <1 μm in diameter, suggesting that intraluminal Aβ deposits locally deform and enlarge NLV structure. **(H)** Image of an individual NLV co-stained with CRANAD-3 and 6E10. (a-b) Normalized fluorescence intensity profiles of CRANAD-3 and 6E10 along the dashed lines in the overlay panel of (H). 6E10-labeled Aβ fibrils adhere to the wall of NLV, and illuminate segments of the vessel, supporting the interpretation that CRANAD-3 reveals NLVs by binding to Aβ species not detectable by 6E10. For all scatter plots, data are presented as mean ± standard deviation (SD), and P values were calculated using nonparametric Mann-Whitney tests. ****P < 0.0001. Scale bars, 10 μm for left panel of (A), 1 μm for right panels of (A); 1 μm for (F) and (H).

**Figure. 7.**
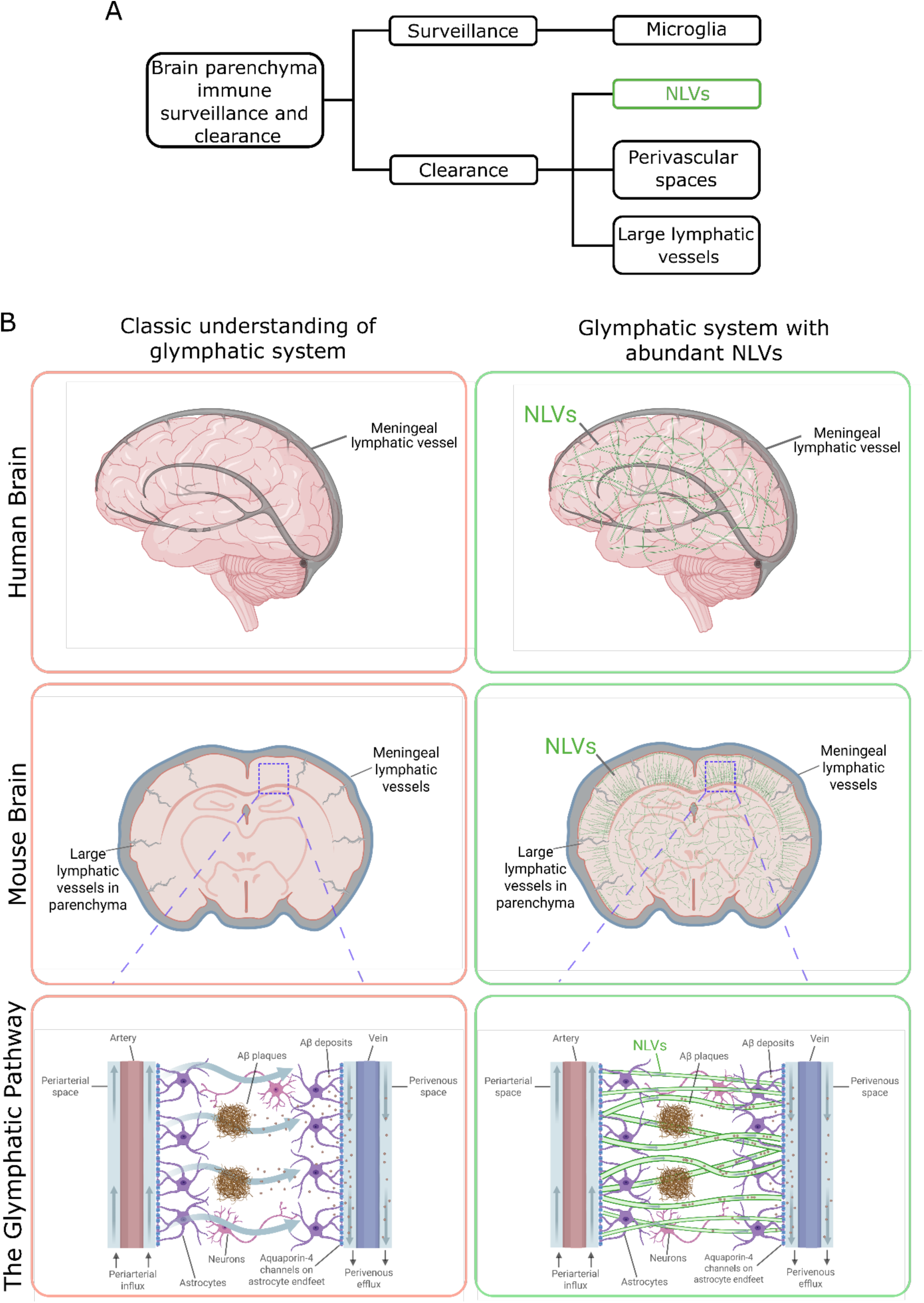
Schematic illustration of immune surveillance and clearance in the brain parenchyma, highlighting the involvement of NLVs and a revised hypothesis of the glymphatic system. (**A**) Illustration of an updated model of immune surveillance and clearance in the brain parenchyma that incorporates NLVs. (**B**) Comparison between the classical view of the glymphatic system and an updated model incorporating the discovery of NLVs. By interacting with arteries and veins via AQP4 channels, NLVs facilitate waste clearance from the brain parenchyma, including small Aβ deposits. Images were created in Biorender.com

### Possible NLV connectivity in parenchyma with ExM and non-ExM

In 2012, Iliff and Nedergaard et al. showed that tracers injected into the cisterna traveled from the PVS of arteries and arterioles to veins ^4^. They proposed a “bulk-flow” mechanism to explain this transport, rather than simple diffusion. However, the nature of this bulk flow and whether it is supported by a structured form remain unclear. To explore this, we investigated the connectivity of NLVs. Visualization of vascular structures required blood vessel markers, but CD31 failed under ExM conditions ^25^. Instead, we used α-SMA, a marker for pericytes on blood vessels ^56^. α-SMA staining with ExM revealed bundles of NLVs coiling around blood vessels (Fig.5A), suggesting a close association between NLVs and the vascular system.

AQP-4, a classical marker for astrocyte ^4,57^, has been widely investigated for its role in transporting CSF in PVS. Abundant AQP-4 expression aligns with the PVS that parallel to blood vessels. To investigate the relationship between NLVs and AQP-4, we used ExM/CRANAD-3 and AQP-4 antibody. We observed that AQP-4 was distributed across the whole tissue, which is expected as astrocytes are widely present in brain tissues. Similar to α-SMA, we found that abundant NLVs coiled around a round hole that is formed by AQP-4 signals (Fig.5B). Moreover, we noticed that AQP-4 puncta were aligned with NLVs (Fig.5C,D), and line plots and quantitative analysis further confirmed this association (Fig.5E). In addition, analysis of single Z-section images revealed a strong association between AQP-4 and CRANAD-3 signals (Fig.5F–I). Together, these findings indicate a close relationship between AQP-4 and NLVs.

Although NLVs can be visualized using both ExM and non-ExM, each approach has limitations. ExM provides rich structural details but can weaken or lose some marker signals, whereas non-ExM preserves markers but lacks detailed resolution. Importantly, we found that the two methods offer complementary advantages. From ExM, we observed certain details of NLV connectivity with blood vessels, but we failed to stain ExM tissue with CD31 ^25^, the most widely used marker for blood capillaries. In this regard, we used non-ExM method to co-stained mouse brain tissue with CD31, AQP-4 and CRANAD-3, to study the connections between blood vessels and NLVs. As reported, we observed that CD31 is surrounded by AQP-4 (Fig.5J,K). While ExM revealed NLVs coiling around areas weakly highlighted by AQP-4, non-ExM provided clearer images confirming this arrangement, with line plots further illustrating the pattern (Fig.5K,L). In addition, we observed putative NLV connections between two AQP4-positive vessels (SI Fig. 5F). Some NLVs appeared to attach to and run parallel to AQP4-positive vessels (SI Fig. 5G), whereas others extended perpendicular to these vessels (SI Fig. 5H). We also observed NLVs tightly coiling around blood vessels (SI Fig. 5I), as well as close contact points between AQP4-positive vessels and NLVs (SI Fig. 5J). However, the precise nature of these interactions requires further confirmation.

We also investigated potential connections between NLVs and classical lymphatic vessels within the brain parenchyma. Using whole-brain clearing and light-sheet imaging, Chang *et al.* reported sparse but classical lymphatic vessels distributed throughout the brain ^18^. Compared with NLVs, these classical lymphatic vessels are considerably larger, with diameters of approximately 5–14 μm. Although we were unable to clearly delineate the contours of classical lymphatic vessels in sectioned brain tissues using VEGFR3, Prox-1, and PDPN staining, hollow structures with diameters of 5–15 μm were detectable with these markers and are likely cross-sections of classical lymphatics. Notably, we observed NLVs coiling around or attaching to the luminal spaces marked by VEGFR3, PDPN, and Prox-1 (SI Fig. 5K–M), suggesting potential connections between NLVs and classical lymphatic vessels.

### Aβ deposits inside NLVs

It has been reported that glymphatics are highly related to the clearance of Aβ in the brain ^4,13,25–28^. Peng et al. reported that the Aβ transportation via glymphatics was significantly reduced in APP/PS1 AD mouse model, compared to WT mice ^28^. Notably, during non-ExM imaging with Aβ antibody (6E10), we observed that Aβ deposits were retained within the NLVs (Fig.6A). Furthermore, Aβ-carrying NLVs exhibited larger diameters (Fig.6B) and higher CRANAD-3 intensities (Fig.6C). We also analyzed the frequency of 6E10 positive dots inside NLVs and found that around 25% of NLVs contained 4-5 dots within 1 micron distance (Fig.6D), and NLVs with more dots showed larger diameters (Fig.6E). Moreover, when we examined single NLVs with Aβ dots at different positions and 6E10 intensities, we found that higher 6E10 intensities were associated with larger diameters (Fig.6F,G). This suggests that Aβ deposits can bulge NLV and make it swollen to larger sizes (more than 1um). In addition, we noticed that Aβ not only could deposit inside NLVs but also could accumulate on the wall of NLVs (Fig.6H). Using another Aβ antibody (4G8), we also observed similar phenomena (SI Fig. 7A,B). Lastly, like non-ExM, we also found that Aβ deposits could be clearly observed inside enlarged NLVs with ExM (SI Fig.7C,D). Taken together, our results suggest that NLVs may play a role in Aβ transport; however, further investigation is required to confirm this function. In addition, it remains unclear whether Aβ aggregation contributes to the enlargement of NLVs

## Discussion

The lymphatic system plays a central role in clearing metabolic waste throughout the body, and a highly efficient lymphatic system is indispensable for maintaining an organ’s health^58^. The brain is among the most metabolically active organs and therefore certainly requires a highly efficient system for waste clearance. In peripheral tissues, this function is supported by a well-organized lymphatic architecture composed of large vessels and abundant capillary lymphatics, which together enable efficient waste clearance. However, such a structured lymphatic system, particularly one comprising abundant capillary lymphatic vessels, has rarely been reported in the brain parenchyma ^17,18,59,60^. It has been proposed that “CSF serves as lymphatics” by exchanging brain ISF along the PVS ^4,9,10^. However, CSF itself lacks an intrinsic structural architecture; instead, it is a collection of liquid solutes in the brain.

In 2012, the glymphatic system was proposed as the lymphatic system in the brain, and large lymphatic vessels were observed, particularly within the meninges and along the cortical surface ^4,9,10^. Nonetheless, abundant capillary lymphatic vessels have not been widely reported within the brain parenchyma. E^18^ven with tissue clearing and advanced light-sheet imaging technologies, the nano-scale structures remain “hidden”. Most of the previous research has focused on large lymphatic vessels and has primarily examined the dura, meningeal, arachnoid regions, while only a few imaging studies have probed lymphatic structures within deep brain parenchyma ^18^. While large lymphatic vessels are comparatively straightforward to detect, the nano-scale vessels are likely to remain “hidden” and “invisible” due to their small dimensions and low SNR in conventional imaging technologies and the limited efficiency of antibody-based labeling strategies.

In this study, we found that the nano-scale structures are widespread across the brain from the meninges/dura to deep brain areas. These NLVs form networks, revealing a previously unrecognized, structured system that may be responsible for the collection and exchange of metabolic waste and other substrates. Such a web-like architecture may provide a clearance mechanism that is substantially more efficient than the simple diffusion previously proposed. Moreover, the potential for structured clearance mediated by NLVs provides a structural basis for the “bulk flow” phenomenon originally proposed by Iliff and Nedergaard in 2012 ^4^. In addition, the NLV system also suggests that CSF–ISF exchange within the parenchyma may also be controlled by NLVs discovered in this letter. Jiang-Xie and Kipnis et al. suggested that large-amplitude, rhythmic ionic waves arising from synchronized neuronal activity in the interstitial fluid could act as a plausible driver for glymphatic flow within the brain parenchyma ^61^. It is possible that the NLV architecture is one of the structural mediators of the high-energy ionic waves in the neural networks, as SMI-positive axons closely intertwine with NLVs in the parenchyma (Fig.4L-N).

In this letter, we describe NLVs as “lymphatic-like,” rather than actual lymphatics, based on two main considerations. First, due to their small sizes, NLVs do not appear to mediate immune cell transport, which is one of the canonical functions of traditional lymphatic vessels; Second, although we demonstrate that canonical lymphatic markers—including LYVE-1, Prox-1, and PDPN—align with NLVs and show strong spatial association, these markers do not delineate clear vessel contours. Notably, under non-ExM conditions, the classical markers LYVE-1, Prox-1, and VEGFR3 could weakly visualize NLV boundaries, although their labeling sensitivity was inferior to that of CRANAD-3. Importantly, both LYVE-1 and CRANAD-3 can be used to visualize the continuous lymphatics and show high co-localization within the meninges, strongly suggesting that the NLVs highlighted by CRANAD-3 are indeed lymphatic vessels. Our study suggests that NLVs are not canonical lymphatics, and new and specific markers beyond LYVE-1, Prox-1, and PDPN will be needed for a deeper understanding of NLVs.

Our discovery of NLVs may also provide certain clear clues about why the brain was deemed immune privilege in the past. According to the canonical view, lymphatic systems integrate two inseparable functions: collection of waste and transportation of immune cells for immune surveillance. In the brain, it is well-established that microglia and astrocytes are responsible for immune surveillance while CSF and ISF serve as the waste carriers; however, what remains unclear is whether the process of waste collection and clearance is mediated by structured systems. Our discovery of NLVs suggests that the lymphatic immune system in brain parenchyma is organized into two components: waste clearance is executed by NLVs, while the immune surveillance is carried out by the residing microglia and astrocytes ^62–65^ (Fig.7A). This two-component mode provides a plausible explanation for why only a few large lymphatic vessels can be found in parenchyma. More importantly, this mode may be leveraged to explain why the brain can efficiently remove waste to meet the high metabolic demands, while microglia maintain immune surveillance ^62–65^.

### Difference between NLVs in parenchyma and meninges

Although all the lymphatic structures in brain parenchyma and meninges are collectively referred to as glymphatics, our findings indicate that their underlying structures can be very different. Meningeal lymphatics, like peripheral lymphatics, exhibit branched structures with heterogeneous diameters, including abundant capillaries. In contrast, within the brain parenchyma, only a few large lymphatic vessels (> 5 μm) can be observed, and abundant capillary lymphatics have not been reported previously. In this letter, we uncovered abundant “hidden” NLVs. The structural differences between these two vessel types likely reflect their distinct functional roles.

Meningeal lymphatics participate in both metabolic waste clearance and immune cell trafficking, whereas parenchymal NLVs seem to serve predominantly as conduits for metabolic waste removal. Notably, meningeal lymphatics seem to be a hybrid of classical lymphatics and NLVs, as we observed both canonical vessels and abundant NLV-like structures within the main trunks and off-trunk regions (SI Fig.8A,B). Upon magnifying these CRANAD-3-stained regions, we detected numerous straight, nano-scale linear structures, likely corresponding to NLVs within the meninges (SI Fig.8A,B). These observations suggest that NLVs may facilitate efficient CSF circulation within the meninges. In addition, we observed canonical lymphatic vessels containing compacted cobblestone-like arrangements with holes measuring 500-1000 nm in diameter (SI Fig.8C,D). However, these cobblestone features are unlikely to be initial lymphatics ^66^, as the holes are arranged regularly within lymphatic vessels. These holes may facilitate fluid communication between different meningeal layers and brain surfaces. Notably, such cobblestone-like vessels were not observed in the brain parenchyma.

Meningeal lymphatics appear to be highly dynamic beyond the trunk regions, as evidenced by lymphangiogenesis features, including spindle-shaped endothelial cells (elongated cells) that are not associated with vessels ^66–68^, but instead appear to be in the process of forming new lymphatic vessels (SI Fig.8E,F).

Beyond meninges, we also observed NLVs in the spinal cord using both ExM and non-ExM approaches, suggesting that NLVs may play an important role in spinal cord function. We further speculate that NLVs are also likely to be present in the retina, although we did not examine retinal tissue in this study. In contrast, no NLVs were observed in peripheral lymph nodes (SI Fig.6).

### Possible connectivity between NLVs and PVS and classical lymphatic vessels

In most peripheral organs, lymphatic vessels typically run parallel to blood vessels as a separate circulatory system. In brain parenchyma, the reported glymphatic vessels also run parallel to blood vessels but as cover-sleeve around the blood vessels, and PVS staining with AQP-4 was often used as the surrogate to highlight the glymphatics ^4,69,70^. However, we observed some NLVs run perpendicular to AQP-4 highlighted glymphatics (SI Fig.5H). Moreover, unlike peripheral lymphatics that have clear branches with different diameters, the NLVs barely have clear branches, and the diameter sizes of NLVs are in a narrow range of 500-1000 nm. However, it remains unclear from our existing data whether the NLVs are branching out from the PVS and classical lymphatic vessels, or largely form a second independent system.

From our observations, the highly abundant NLVs tend to bundle or coil around AQP-4-positive vessels, and these bundled NLVs likely contribute to the formation of PVS. Such bundling provides a plausible explanation for why cisternal tracers accumulate along the perivascular rim. In contrast, non-bundled NLVs yield minimal detectable signal following cisternal tracer injection, likely because the dispersed NLV density dilutes the tracer and causes the signal to be obscured by background noise. In vitro brain tissue immuno-histological staining often showed that AQP-4 and CD31 are very closely attached to each other while AQP-4 is on the abluminal side, and the distance between these two is often less than 2 microns ^71,72^. However, in vivo two-photon imaging in Iliff and Nedergaard’s study showed that the sizes of PVS are around 20 microns ^4^. In this regard, NLVs may be used to interpret this discrepancy, as Fig.5K showed that the bundled NLVs form a void space around the vessels visualized by AQP-4/CD31. Regarding the potential connections between NLVs and classical lymphatics in the parenchyma, although we were unable to clearly delineate their contours using Prox-1, PDPN, and VEGFR3 staining in sectioned brain slides, possible NLV connections were observed around the hollow structures that were faintly highlighted by these markers.

### Potential NLV functions and its relations with diseases

In this letter, we have not conducted experiments to investigate NLV functions; the following discussion is therefore speculative. Iliff and Nedergaard et al. showed that once the tracer is injected into the cisterna magna, it rapidly enters the PVS but then slows markedly, suggesting the presence of physical barriers along this route ^4^. It is plausible that astrocytic endfeet together with NLVs contribute to these barriers, restricting tracer movement. Evidence for size selective exclusion also exists: large molecules such as FITC–Dextran 2000 (2,000 kDa; ∼322 nm) rarely reach the venous side following cisternal injection ^4^, indicating that such barriers prevent the passage of macromolecules. Additionally, the absence of Gd-MRI contrast leakage into the parenchyma after cisternal administration further supports the idea that these barriers restrict even certain small molecules from entering the brain parenchyma ^9,73–75^.

Our current understanding is that glymphatics mediate waste clearance by transporting metabolites out of the brain, a view strongly supported by numerous studies. In this work, we observed Aβ deposits within NLVs, further indicating that NLVs may participate in this transport process and are integral components of the broader clearance network.

Our observations indicate that NLVs are tightly regulated, and they play important roles in both normal physiology and disease. In AD mouse brains, several notable alterations emerge relative to wild-type controls. NLVs often appear fragmented, and regions immediately surrounding amyloid plaques show conspicuous NLV-free zones. In some cases, NLVs adjacent to plaques are displaced and compressed into dense bundles rather than maintaining an even distribution. This pattern suggests that NLVs are structurally flexible—likely less rigid than blood vessels or capillaries—and can be physically shifted by local pathological structures. Because of their high abundance and flexibility, NLVs may also be resilient and re-generated ^16,76,77^; minor damage to individual nano- or micro-vessels could be readily compensated by neighboring vessels.

The initial goal of our study was to determine whether Aβ species extend beyond plaque-enriched regions. Using our fluorescence probe CRANAD-3, we visualized Aβ and confirmed that its distribution is not restricted to plaques or their immediate rims but instead spans across broad brain areas. As we continued our investigation, the abundance and web-like organization of NLVs became increasingly apparent. This raises an important question: how does the NLV network contribute to AD pathology, and why certain regions might be more vulnerable than others. Additionally, we observed that some NLVs traverse multiple cortical layers. Such long-range trajectories may provide an additional structural pathway for the spread of α-synuclein and other misfolded proteins, complementing the well-established axonal propagation routes ^78–81^.

Our study with CRANAD-3 probably suggests that Aβ species are broadly distributed across multiple brain regions, the meninges, and the spinal cord. Although the dominant Aβ species in NLVs remain uncertain, these findings raise the possibility that Aβs may be required for maintaining NLV structure and function. Nonetheless, it is clear that excessive accumulation and deposition of Aβ within NLVs are detrimental to brain function.

### CRANAD-3 specificity

During our non-ExM imaging of whole-brain sections, we observed CRANAD-3 fluorescence in white matter regions of the corpus callosum and in fiber bundles within the putamen. This signal may arise from two possibilities: (a) CRANAD-3 is not fully specific for Aβ and may exhibit some nonspecific affinity for myelin; or (b) CRANAD-3 binds to Aβ located within internodal periaxonal spaces and paranodal/juxtaparanodal channels, which were recently reported ^82^. Notably, we also detected similar spiral patterns of Aβ using 6E10 antibody staining in neuronal axons as reported ^82^ (SI Fig.8G,H).

Our results indicate that CRANAD-3 provides high-quality visualization of NLV contours, and its primary labeling is likely driven by its affinity for Aβ. However, the fluorescence signal within NLVs may not exclusively reflect Aβ binding; CRANAD-3 may also interact with other proteins, such as the β-amyloid precursor protein (APP) in endothelial cells ^83^.

### Nano-scale vascular and tubular structures in the brain

Recent advances in imaging technologies have revealed several remarkable nano-scale structures in the brain. Rustom et al. first identified nanotubular networks that mediate intercellular communication by transporting organelles between cells^84^. More recently, Chang and Kwon et al. described a dendritic nanotubular (DNT) network that enables communication among brain cells ^85^. Using super-resolution microscopy in dissociated cortical neurons, they showed that DNTs are rich in actin, support long-range calcium (Ca²□) propagation, and can transport Aβ to neighboring neurons. Importantly, disruptions in the DNT network arise early in disease progression, preceding the formation of amyloid plaques. Rakotobe et al. reported tunneling nanotube (TNT)–like connections in the developing cerebellum, distinct from cytokinetic and intercellular bridges^86^. These TNT structures also participate in intercellular communication. Additionally, Alarcón-Martínez and Di Polo et al. demonstrated that inter-pericyte TNTs regulate neurovascular coupling ^87,88^. Scheiblich, Heneka, and others further showed that in postmortem brain tissues from patients with Lewy body disease or multiple system atrophy, α-synuclein–laden microglia extend TNTs to less-burdened neighboring microglia to enhance pathogenic α-synuclein clearance ^89^.

The NLVs described in this letter are fundamentally distinct from TNT and DNT networks, as they operate beyond intercellular signaling and likely plays broader roles in molecular transport across the brain. Nonetheless, all these nano-scale networks share a common feature: they offer far more efficient routes for biomolecule movement than simple diffusion, facilitating communication and transport across cells and brain regions. With continued progress in biomedical imaging, additional nano-scale structural networks are likely to be discovered, further deepening our understanding of brain organization and function.

### Potential role of NLVs in diffusion tensor imaging (DTI) and fMRI

Diffusion tensor imaging (DTI) and diffusion tractography assess the directional diffusion of water molecules in tissue, based on the assumption that diffusion anisotropy reflects underlying white matter fiber orientation ^90,91^. Interestingly, our non-ExM imaging of NLVs revealed orientation patterns in the cortex similar to fibers labeled by SMI-31 antibodies. Like fiber tracks, water movement within NLVs is also diffusion-restricted. In this regard, the NLV network may help address some of the unresolved questions in DTI interpretation. Furthermore, vessel wall motion—including arterial pulsations and slow vasomotion—generates pressure fluctuations and fluid dynamics in perivascular spaces, which are critical for CSF inflow and waste clearance ^9^. The NLV network may serve as a structural basis for studying such oscillatory flows. In addition, slow neural activity oscillations (<1 Hz; often 0.01–0.1 Hz) are a hallmark of resting-state brain activity and contribute substantially to functional connectivity measured by fMRI ^92^. Therefore, the NLV network may provide valuable insights for interpreting fMRI signals.

### Why NLVs have been missed before

Why has such an important organized system remained undetected for so long? Several important factors likely contributed. First, although LYVE-1 is considered the gold-standard marker for lymphatic vessels, it reliably labels only larger glymphatic structures. In brain parenchyma, LYVE-1 staining frequently appears patchy, discontinuous, or even punctate ^18^, making it unsurprising that nano-scale glymphatics cannot be visualized either by LYVE-1 immunostaining or in LYVE-1–GFP reporter mice. Similarly, other commonly used lymphatic markers—including Prox-1, PDPN, and VEGFR3—lack the sensitivity or spatial resolution needed to delineate nano-scale lymphatic structures. Second, major imaging modalities such as MRI, PET, and CT do not reach the spatial resolutions required to resolve vessels at nanometer to sub-micrometer scales. Although transmission electron microscopy (TEM) can reach such resolution, it is limited to extremely small fields of view, and reconstruction of extended vascular networks requires labor-intensive stitching. Moreover, TEM often requires specialized sample preparation and staining to generate sufficient contrast. Third, although our findings demonstrate that fluorescent Aβ probes like CRANAD-3 can reveal NLVs, conventional brain sections typically yield low signal-to-noise ratios under fluorescence microscopy. Such low SNR compromises not only standard high-resolution imaging but also super-resolution techniques, leading to failure in visualizing NLV contours. Aβ immunostaining combined with ExM has the potential to reveal NLVs, as linearly aligned nano-clusters have been reported, yet the continuous nano-scale vessel morphology remained largely unresolved in those studies ^22^. Lastly, the brain parenchyma differs fundamentally from peripheral tissues: under physiological conditions it contains very few T cells, and its lymphatic-like vessels are not responsible for routine immune-cell transport. This suggests that capillary-scale lymphatics in the parenchyma may have distinct structural properties compared to peripheral lymphatic capillaries. Nevertheless, it has been difficult to envision that their morphology diverges so sharply, which likely contributed to their being overlooked. Taken together, limitations in molecular markers, imaging resolution, fluorescence signal quality, and assumptions about parenchymal lymphatic morphology collectively explain why NLVs remained hidden until now.

### Prospectives and summary

Using the Aβ[binding probe CRANAD-3, we successfully revealed previously hidden NLVs in the brain, suggesting that Aβ species may serve as provisional markers for these structures. Nevertheless, more specific and definitive molecular markers will be required to fully understand NLVs. Although our ExM and non-ExM imaging provided extensive characterization of NLV structures, additional approaches—such as tissue clearing combined with light-sheet microscopy—may further enable whole-brain mapping and reconstruction of the NLV network. Many important questions remain unanswered, including how NLVs interface with larger lymphatic pathways in the parenchyma and meninges, and whether NLVs possess valve-like structures.

Our discovery of NLVs in the brain opens a new avenue for neuroscience and related fields. Together with perivascular spaces and meningeal lymphatics, the NLV system may contribute to a more complete picture of the glymphatic system. Our finding provides a crucial missing piece in understanding glymphatic structures and functions, and continued investigation of the NLV system will undoubtedly deepen our insight into this system.

## Acknowledgments

This work was supported by NIH grants R01AG055413, R01AG085562, R21AG080222, R21AG078749, and S10OD028609 awards (C. R.). The authors also wish to thank Dr. Bruce R. Rosen, MD, PhD, for his valuable feedback on this manuscript. The authors would also like to thank Ms. Alice Li and Ms. Amy Wang for their assistance in preparing experimental materials.

## Notes

### Competing Interest Statement

The authors have declared no competing interest.

### Summary of Updates

The authors have replaced the abstract with a revised version to improve clarity. In addition, we also changed the author list.

