## Supplementary material for "Discovery of Abundant Nano-scale Lymphatic-like Vessels in Brains": Methods and SI Figures

### Materials and Methods

**Mouse brain tissue preparation**. All animal experiments were performed in accordance with protocols approved by the Institutional Animal Care and Use Committee (IACUC) of Massachusetts General Hospital and complied the U.S. Public Health Service Policy on Humane Care and Use of Laboratory Animals. Female wild-type mice (C57BL/6J, 2 or 8.5 months of age), and 5xFAD mice (B6SJL-Tg, 8.5 months of age) were purchased from The Jackson Laboratory. To harvest mouse brain tissue and meninges, mice were first deeply anesthetized with 3% isoflurane. They were next transcardially perfused with ice-cold 10 ml 1X phosphate-buffered saline (PBS), followed by 10 ml of 4% paraformaldehyde (PFA) in 1X PBS at room temperature (RT). Mouse brain tissue and meninges were carefully collected and soaked in 10 ml of 4% PFA in 1X PBS at 4°C for 24 hours. Fixed tissues were briefly washed twice with 1X PBS and dehydrated in 10 ml of 20% sucrose in 1X PBS supplied with 100 mM glycine at 4°C for 24 hours, followed by 30% sucrose incubation for another 24 hours at 4°C. After dehydration, meningeal tissues were kept in 1X PBS for 2 days at 4°C to facilitate the separation step. Brain tissues were immediately embedded and frozen in O.C.T. compound using acetone solution supplied with dry ice. Brain slices (30 μm) were cut on a cryostat (Leica) and stored in cryoprotectant at -20°C until staining.

**Immunostaining.** Mouse brain slices were rinsed 3 times with 1X PBS, and incubated in blocking solution (0.3% TrigonX-100, 5% normal goat serum (NGS) in 1X PBS) for 2 h at RT. Tissues were then incubated with primary antibodies (see Methods Primary antibody list) in blocking solution overnight at 4°C. On the following day, tissues were washed in washing solution (0.3% Triton X-100 in 1X PBS) 3 times for 5 minutes each at RT, followed by secondary antibody (see Methods Secondary antibody list) incubation in blocking solution for 2 h at RT. During the last 15 minutes of secondary antibody incubation, CRANAD-3 was diluted inside to a final concentration of 10 μM. All tissues were then washed 3 times with washing solution and 3 times with 1X PBS. Tissues were either mounted on a glass slide with Prolong Diamond Antifade Mountant (Thermo Fisher Scientific) and coverslips, or briefly air dried on glass bottom transparent Petri dishes (Ted Pella Inc) and imaged in 1X PBS.

**Tissue expansion and staining**. Anchoring and gelation steps were performed according to the protocol (Valdes et al. 2024). Briefly, mouse brain slices were first briefly rinsed 3 times with 1X PBS, followed by overnight incubation with anchoring agent Acryloyl-X solution (AcX) at a final concentration of 0.1mg/ml in 1X PBS supplied with 0.5% Triton X-100 at 4°C. On the next day, the anchored tissues were washed 2 times with 1X PBS, 10 minutes each at RT. The tissues were next incubated with gelation solution by mixing following chemicals: 0.01% 4-hydroxy-2,2,6,6-tetramethylpiperidin-1-oxyl (4-HT), 0.2% (w/v) tetramethylethylenediamine (TEMED), and 0.2% (w/v) ammonium persulfate (APS) in monomer solution (2 M sodium chloride (NaCl), 8.625% (w/v) sodium acrylate, 2.5% (w/v) acrylamide, and 0.10% (w/v) *N*,*N*′-methylenebisacrylamide in 1X PBS). During the gelation step, on a rectangular coverslip (24 mm by 60 mm), one No. 1.5 square coverslip (22 mm by 22 mm) was cut into two halves as two spacers and placed 15mm apart, and 80 ul of gelation solution were applied between the two spacers. After transferring the brain tissue to this 80 ul gelation solution, another square coverslip was carefully placed on top of the two spacers to enclose the chamber so that the tissue was lying flat inside. The gelation chamber was maintained in a humidified environment and incubated for 0.5 h at 4°C, followed by 37°C incubation for 2.5 h.

Upon completion of the gelation step, all coverslips were carefully removed. A razor blade was used to remove the excess gel around the tissue. The digestion buffer was prepared by mixing the following chemicals: 20% (w/v) SDS, 25 mM EDTA (pH 8.0), 50 mM Tris (pH 8.0), and 0.5% Triton X-100. The pH of the digestion buffer was adjusted to 8.0 with hydrochloric acid, followed by adding 100 mM β-mercaptoethanol (BME). The trimmed tissues were incubated in the digestion buffer for 0.5 h at 37°C and 1 h at 121°C. After digestion, tissues were allowed to cool to RT for around 0.5 h and they were washed multiple times with 1X PBS. The digested tissue, which has already reached ~2x expansion, can be stored in 1X PBS at 4°C fridge for several weeks.

The digested tissues were then performed with immunostaining if needed (without staining CRANAD3 in the final step) as explained above. Tissues were then submerged in distilled water to allow expansion and replaced with fresh water every 10 minutes for around 3-5 times until they reach a final 4X expansion. The expanded gels were carefully transferred to Poly-D-Lysine (PDL)-coated glass bottom petri dishes and stayed for around 0.5 h to allow full binding. Tissues were next stained with 10μM of CRANAD3 in distilled water for 10 minutes in dark at RT, followed by 6 washes with distilled water.

**Image Processing and Analysis**. Imaging was performed using either Leica Stellaris confocal and two-photon (2P) microscope at Harvard Medical School, or Nikon AXR confocal microscope at Massachusetts General Hospital. All images were analyzed with ImageJ (NIH, Bethesda, MD) software. To calculate normalized fluorescence intensity, raw fluorescence signals were first subtracted by the minimum value and then divided by the difference between the maximum and minimum values.

**Statistical Analysis.** All values are reported as mean ± SD and statistical analyses were performed and reported using GraphPad Prism 10 (GraphPad Software, Inc., San Diego, CA). Statistical significance was assessed with nonparametric Mann–Whitney tests. All results were considered significant for $P < 0.05$. *$P < 0.05$, **$P < 0.01$, ***$P < 0.001$, and ****$P < 0.0001$; n.s. indicates no significant difference.

Primary antibody list:

| Target | Host | Catalog Number | Vendor |
|---|---|---|---|
| LYVE-1 | Rabbit | ab14917 | Abcam |
| Prox-1 | Rabbit | 925201 | BioLegend |
| PDPN | Rat | ab256559 | Abcam |
| VEGFR3 | Rat | 14-5988-82 | ThermoFisher |
| CD31 | Rat | 550274 | BD Bioscience |
| AQP-4 | Rabbit | HPA014784-100UL | Millipore Sigma |
| 6E10 | Mouse | 803001 | BioLegend |

| | | | |
|---|---|---|---|
| 12F4 | Mouse | 805501 | BioLegend |
| 4G8 | Mouse | 800701 | BioLegend |
| Ankyrin-G | Mouse | 75-146 | antibodiesinc |
| MAP2 | Chicken | ab5392 | Abcam |
| Collagen-I | Rabbit | ab270993 | Abcam |
| SMA | Mouse | NBP2-33006-0.1mg | Novus Biological |
| SMI | Chicken | ab4680 | Abcam |

Secondary antibody list:

| Conjugation | Host | Species Reactivity | Catalog Number | Vendor |
|---|---|---|---|---|
| Alexa Fluor™ 488 | Goat | Rabbit | A-11008 | ThermoFisher |
| Alexa Fluor™ 488 | Goat | Rat | A-11006 | ThermoFisher |
| Alexa Fluor™ 488 | Goat | Mouse | ab150117 | Abcam |
| Alexa Fluor™ 488 | Goat | Chicken | ab150169 | Abcam |
| Alexa Fluor™ 405 | Goat | Rabbit | A48254 | ThermoFisher |
| Alexa Fluor™ 405 | Goat | Rat | ab175671 | Abcam |
| Alexa Fluor™ 405 | Goat | Mouse | A-31553 | ThermoFisher |

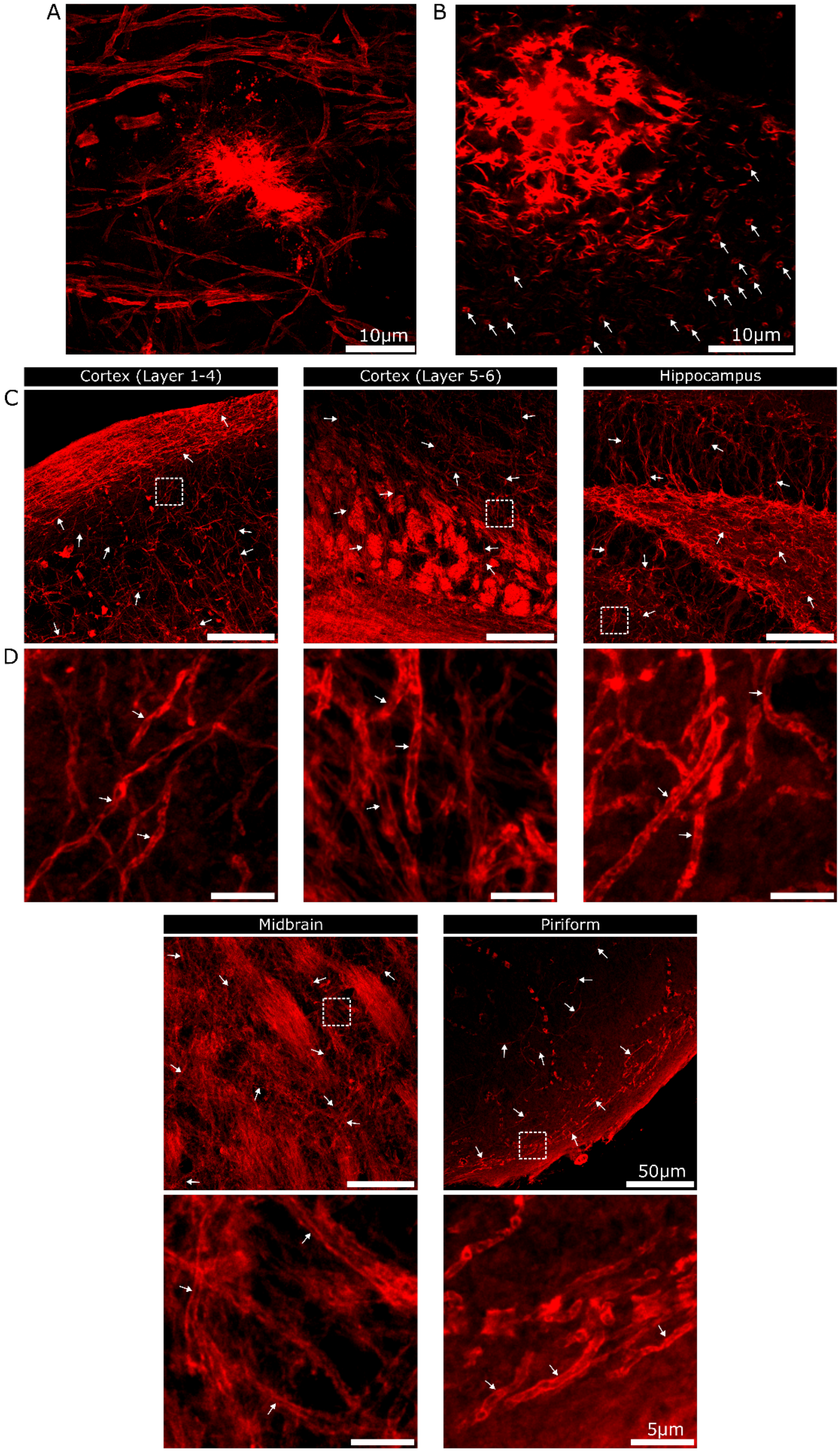

A
B
10µm
10µm
Cortex (Layer 1-4)
Cortex (Layer 5-6)
Hippocampus
C
D
Midbrain
Piriform
50µm
5µm

**SI Fig.1. NLVs ubiquitously exist in mouse brain.**
**(A)** Representative ExM image of an Aβ plaque and several NLVs surrounding it.
**(B)** Cross-sectional view of multiple NLVs around an Aβ plaque. White arrows mark the cross-sections of NLVs with hole/hollow features.
**(C)** ExM images demonstrating the presence of NLV in different regions of a mouse brain, including different layers of cortex, hippocampus, middle brain and piriform cortex. White arrows point to individual NLVs.
**(D)** Magnified views of the boxed regions in (C). White arrows point out some NLVs in each region.

Scale bars, 10 μm for (A) and (B); 50 μm for (C); 5 μm for (D).

SI Movie 1: Three-dimensional movie of NLVs in an ExM mouse brain section stained with CRANAD-3.

SI Movie 2: Z-stack movie of NLVs in an ExM mouse brain section stained with CRANAD-3.

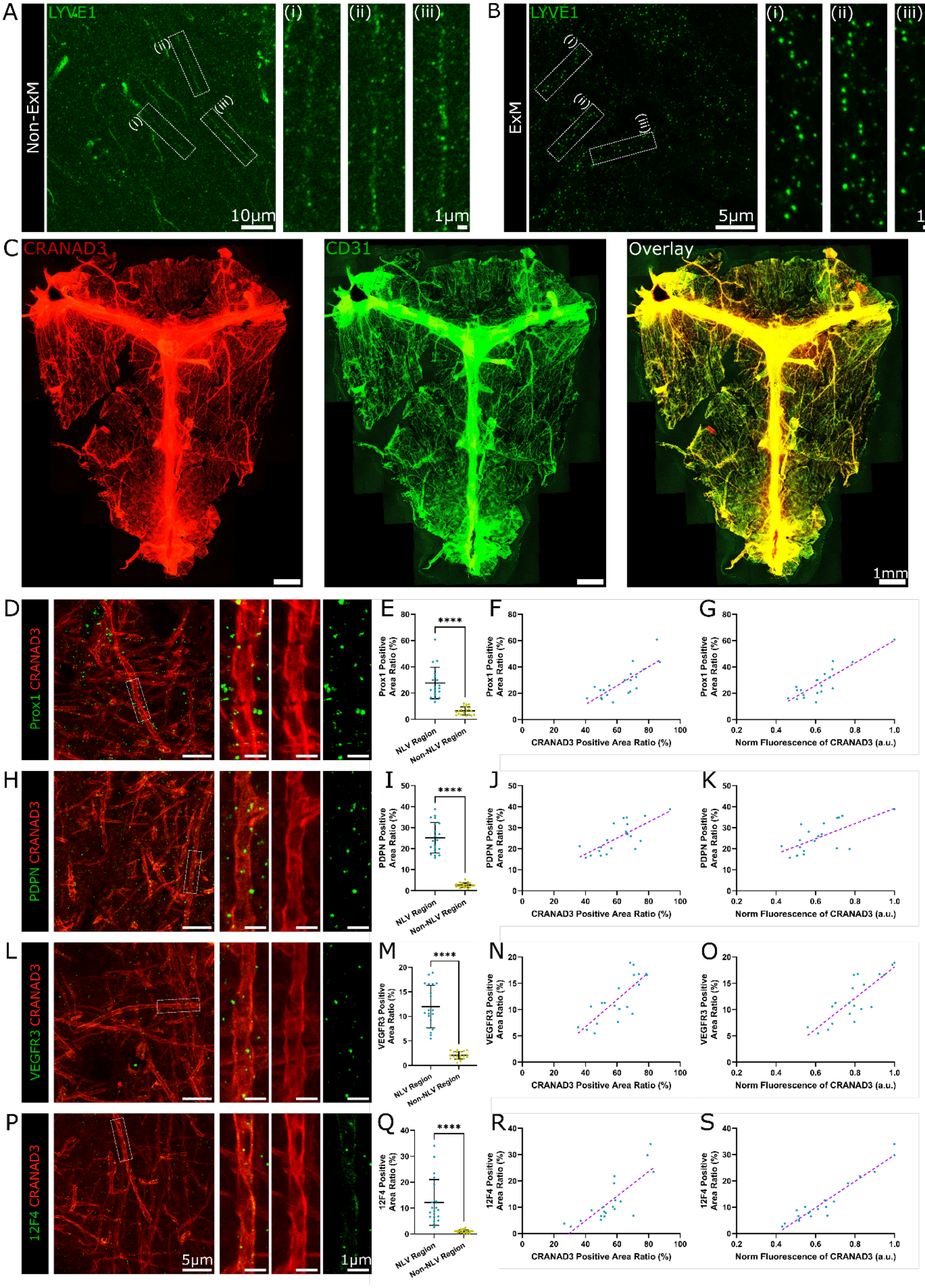

A
Non-ExM
LYVE1
(i)
(ii)
(iii)
10μm
1μm
B
ExM
LYVE1
5μm
1μm
C
CRANAD3
CD31
Overlay
1mm
D
Prox1 CRANAD3
E
Prox1 Positive Area Ratio (%)
NLV Region
Non-NLV Region
****
F
CRANAD3 Positive Area Ratio (%)
G
Norm Fluorescence of CRANAD3 (a.u.)
H
PDPN CRANAD3
I
PDPN Positive Area Ratio (%)
J
K
L
VEGFR3 CRANAD3
M
VEGFR3 Positive Area Ratio (%)
N
O
P
12F4 CRANAD3
Q
12F4 Positive Area Ratio (%)
R
S
5μm
1μm

**SI Fig.2. Additional Validation of lymphatic-like features of NLVs in WT meninges and parenchyma of 5xFAD mouse brain.**

**(A)** Representative non-ExM image of mouse brain slice stained with LYVE-1. (A) (i-iii) Magnified views of the boxed regions in the left panel of (A).

**(B)** ExM image of mouse brain slice stained with LYVE-1. (B) (i-iii) Magnified views of the boxed regions in the left panel of (B).

**(C)** Non-ExM images of whole-mounted WT mouse meninges stained with CRANAD-3 (red) and CD31 (green).

**(D, H, L, P)** ExM image of 5xFAD mouse brain section co-stained with CRANAD-3 and Prox1 (D), PDPN (H), VEGFR3 (L), or 12F4 (P). Right panels show magnified views of the boxed regions in the corresponding left panels.

**(E, I, M, Q)** Quantification of Prox1 (E), PDPN (I), VEGFR3 (M), or 12F4 (Q) normalized fluorescence intensities at NLV areas (n = 20) and non-NLV areas (n = 20) ($P < 0.0001$).

**(F, J, N, R)** Relationship between Prox1- (F), PDPN- (J), VEGFR3- (N), or 12F4- (R) positive area ratios and their correlation with CRANAD-3-positive areas at NLV regions (n = 20) [$R^2$ = 0.6078 for (F); 0.5407 for (J); 0.6182 for (N); 0.6408 for (R); linear regression fitting].

**(G, K, O, S)** Relationship between Prox1- (G), PDPN- (K), VEGFR3- (O), or 12F4- (S) positive area ratios and normalized fluorescence intensities of CRANAD-3 at NLV regions (n = 20) [$R^2$ = 0.7515 for (G); 0.4380 for (K); 0.6699 for (O); 0.9285 for (S); linear regression fitting].

For all scatter plots, data are presented as mean ± standard deviation (SD), and P values were calculated using nonparametric Mann-Whitney tests. ****$P < 0.0001$.

Scale bars, 10 µm for left panel of (A), 1 µm for right panels of (A); 5 µm for left panel of (B), 1 µm for right panels of (B); 1mm for (C); 5 µm for left panels of (D), (H), (L), (P), 1 µm for right panels of (D), (H), (L), (P).

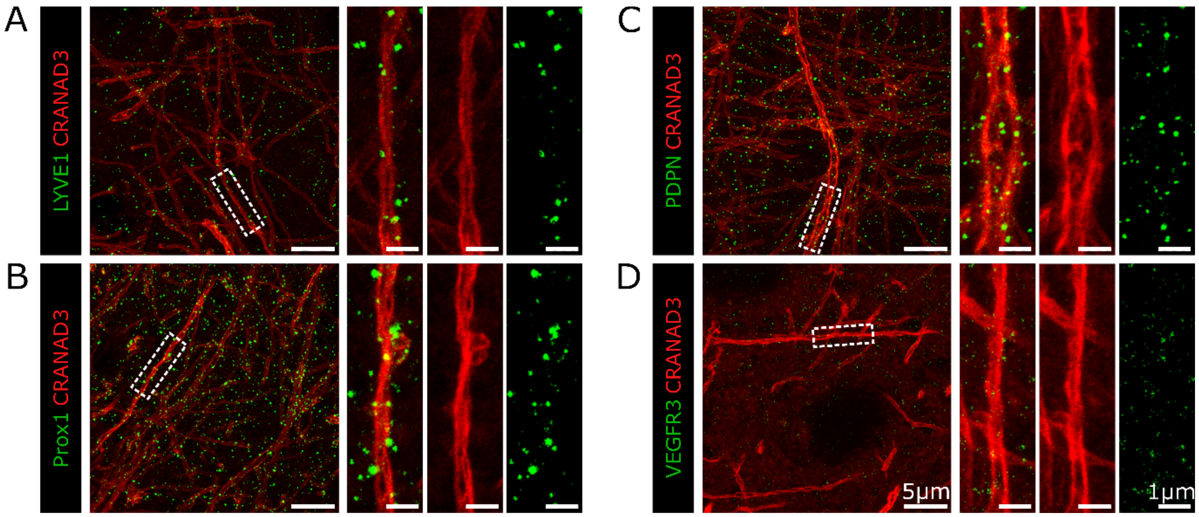


**SI Fig.3. NLVs validation in the parenchyma of WT mouse brain with lymphatic markers.**

**(A-D)** Representative ExM images of WT mouse brain sections co-stained with CRANAD-3 and four lymphatic markers: LYVE-1 (A), Prox1 (B), PDPN (C), and VEGFR3 (D). Right panels show magnified views of the boxed regions in the corresponding left panels.

Scale bars, 5 μm for left panels of (A-D), 1 μm for right panels of (A-D).

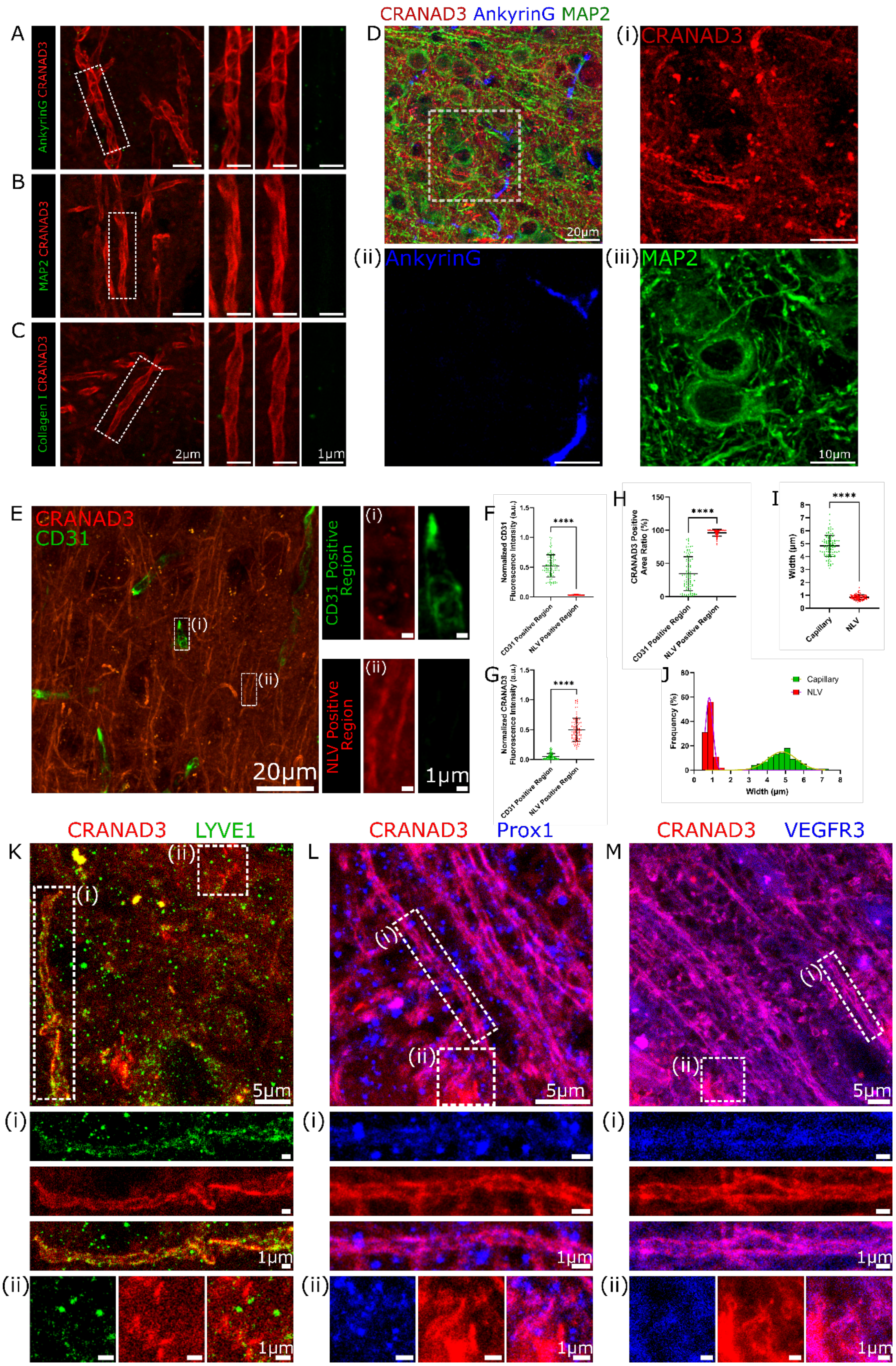

CRANAD3 AnkyrinG MAP2
A
AnkyrinG CRANAD3
B
MAP2 CRANAD3
C
Collagen I CRANAD3
2µm
1µm
D
20µm
(i) CRANAD3
(ii) AnkyrinG
(iii) MAP2
10µm
E
CRANAD3
CD31
(i)
(ii)
20µm
CD31 Positive Region
NLV Positive Region
1µm
F
Normalized CD31 Fluorescence Intensity (a.u.)
****
CD31 Positive Region
NLV Positive Region
G
Normalized CRANAD3 Fluorescence Intensity (a.u.)
H
CRANAD3 Positive Area Ratio (%)
I
Width (µm)
Capillary
NLV
J
Frequency (%)
Width (µm)
Capillary
NLV
K
CRANAD3 LYVE1
L
CRANAD3 Prox1
M
CRANAD3 VEGFR3
5µm

**SI Fig.4. Validation of NLVs that are distinct from neuronal structures and blood vessels.**
**(A-C)** Representative ExM images of WT mouse brain sections co-stained with CRANAD-3 and neuronal markers AnkyrinG (A), MAP2 (B), or the collagen maker Collagen I (C). Right panels show magnified views of the boxed regions in the corresponding left panels.
**(D)** Non-ExM co-staining with CRANAD-3, AnkyrinG, and MAP2. Boxed region is magnified and shown in (i-iii), demonstrating no signal overlap between CRANAD3 with AnkyrinG or MAP2.
**(E)** Non-ExM co-stained with CRANAD-3 with CD31. (i-ii) Magnified images of the boxed regions in (E), indicating distinct spatial separation of CD31-positive blood vessels and CRANAD-3-labeled NLVs.
**(F-G)** Quantitative analysis of normalized CD31 (F) and CRANAD-3 (G) fluorescence intensities in CD31-positive and CRANAD-3-positive (NLV) regions (n = 100 per group; $P < 0.0001$ for both analyses).
**(H)** Comparison of CRANAD-3-positive area percentage between blood capillaries and NLVs (n = 100 per group; $P < 0.0001$).
**(I)** Comparison of NLV widths between blood capillaries and NLVs, showing NLVs are significantly thinner than capillaries (n = 100 per group; $P < 0.0001$).
**(J)** Percentage histogram of blood vessel and NLV widths (n = 100 per group; $R^2 = 0.9567$ for capillary group; $R^2 = 0.9992$ for NLV group; Gaussian nonlinear regression).
**(K-M)** Non-ExM images co-stained with CRANAD-3 and lymphatic markers: LYVE1 (K), Prox1 (L), and VEGFR3 (M). Boxed region (i) and (ii) are shown in the lower panels of each image. For each lymphatic marker, region (i) suggests a moderate to weak overlap with CRANAD-3 signal; region (ii) shows that the signals from lymphatic markers are not caused by the bleed-through or leakage from CRANAD-3 signal.

For all scatter plots, data are presented as mean ± standard deviation (SD), and P values were calculated using nonparametric Mann-Whitney tests. ****$P < 0.0001$.

Scale bars, 2 µm for left panels of (A-C), 1 µm for right panels of (A-C); 20 µm for (D), 10 µm for (i-iii); 20 µm for left panels of (E), 1 µm for right panels of (E); 5 µm for top panels of (K-M) , 1 µm for (i) and (ii) of (K-M).

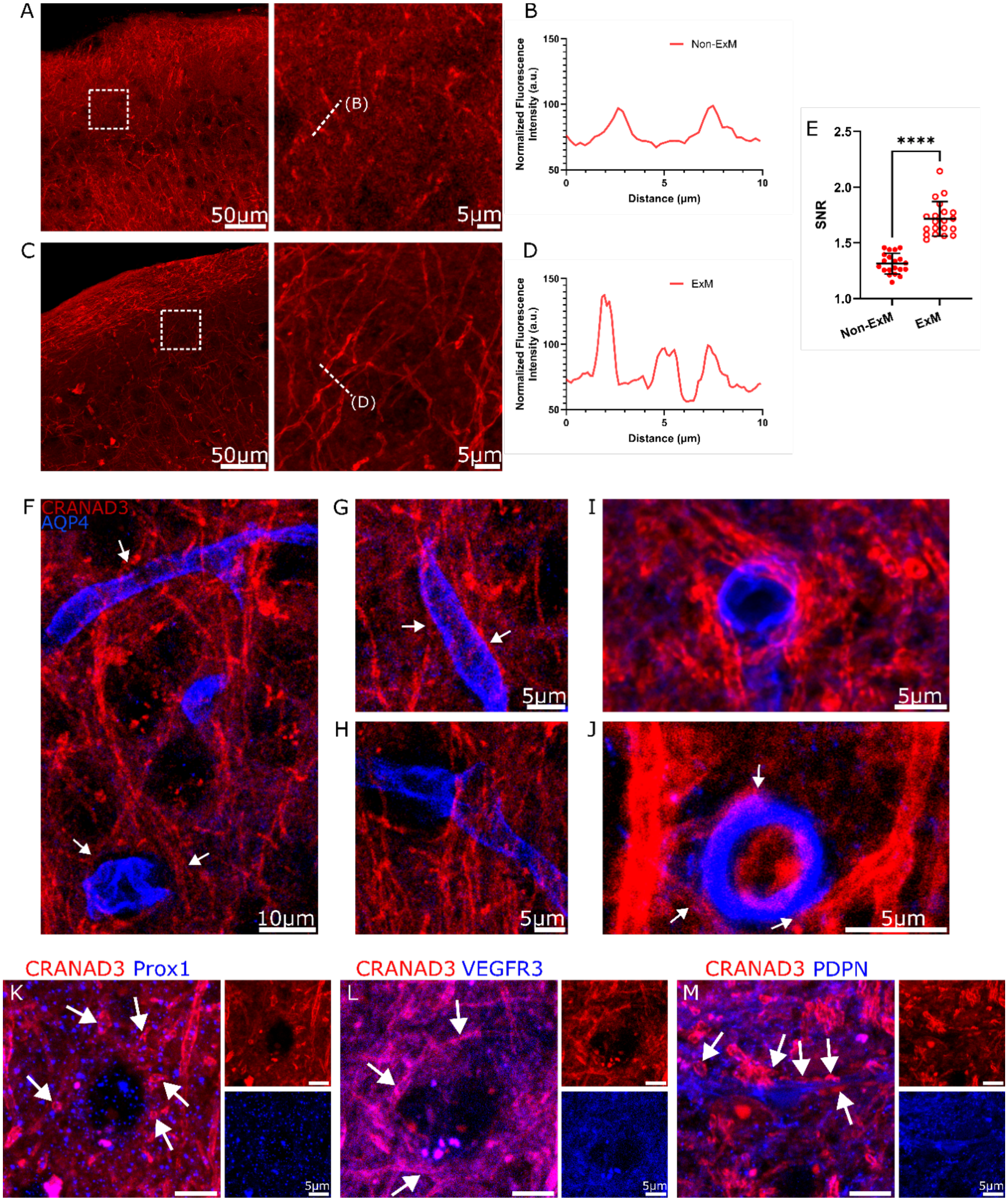


**SI Fig.5. Comparison of non-expanded and expanded imaging of NLVs.**

**(A)** Non-ExM image of mouse brain cortex motor area stained with CRANAD-3. Magnified view of the boxed region is shown on the right.

**(B)** Normalized fluorescence intensity profile of CRANAD-3 along the dashed line in (A).

**(C)** ExM image of mouse brain cortex motor area stained with CRANAD-3. Right panel presents the zoomed-in view of the boxed region.

**(D)** Normalized fluorescence intensity profile of CRANAD-3 along the dashed line in (C). Noise levels were proportionally adjusted between (B) and (D) to achieve comparable baselines for visualization.
**(E)** Comparison of SNR between non-ExM and ExM images acquired at layer 2. SNR is significantly higher in the ExM group. ($n = 20$ per group; $P < 0.0001$).
**(F-J)** Non-ExM imaging with CRANAD-3 and AQP-4 staining in cortex areas to show possible NLV connections between blood vessels and PVS. (F) Possible connections between two blood vessels highlighted by AQP-4 (white arrows). (G) Representative NLVs attached to blood vessels. (H) NLVs run nearly perpendicularly to blood vessels. (I) NLVs tightly wrap around blood vessels. (J) High resolution image shows the possible connection points of three NLVs (white arrows) with artery highlighted by APQ-4.
**(K-M)** Non-ExM imaging with CRANAD-3 and lymphatic markers, including Prox1 (K), VEGFR3 (L), and PDPN (M). White arrows show that several NLVs wrap around classical lymphatic vessels.

For all scatter plots, data are presented as mean ± standard deviation (SD), and P values were calculated using nonparametric Mann-Whitney tests. ****$P < 0.0001$.

Scale bars, 50 μm for left panels of (A) and (C), 5 μm for right panels of (A) and (C); 10 μm for (F); 5 μm for (G), (H), (I), and (J); 5 μm for (K), (L), and (M).

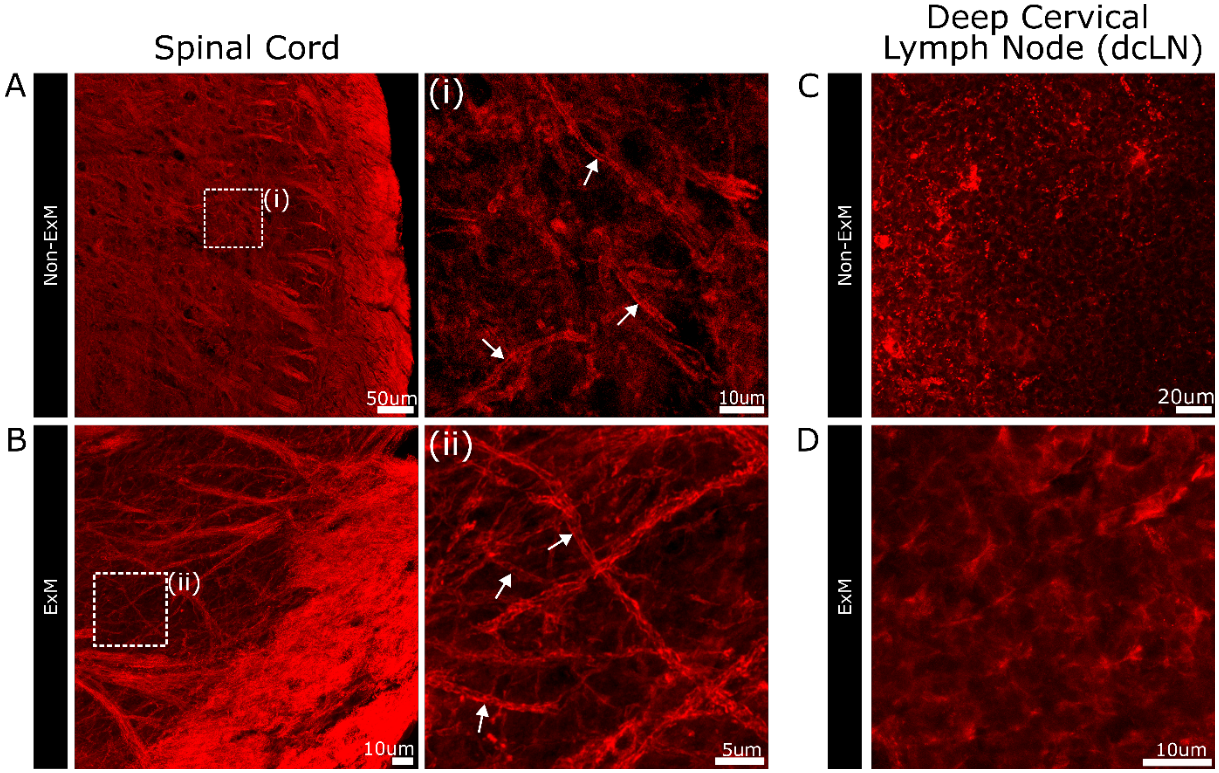


**SI Fig.6. NLVs present in the spinal cord of WT mouse, but not in deep cervical lymph node.**

**(A-B)** Non-ExM (A) and ExM (B) images of WT mouse spinal cord sections stained with CRANAD-3. Magnified views of individual NLVs from the boxed regions in (A) and (B) are shown in (i) and (ii), respectively. White arrow mark individual NLVs.

**(C-D)** Non-ExM (C) and ExM (D) images of deep cervical lymph node (dcLN) sections stained with CRANAD-3, showing no detectable NLV-like structures.

Scale bars, 50 µm for (A), 10 µm for (i); 10 µm for (B), 5 µm for (ii); 20 µm for (C); 10 µm for (D).

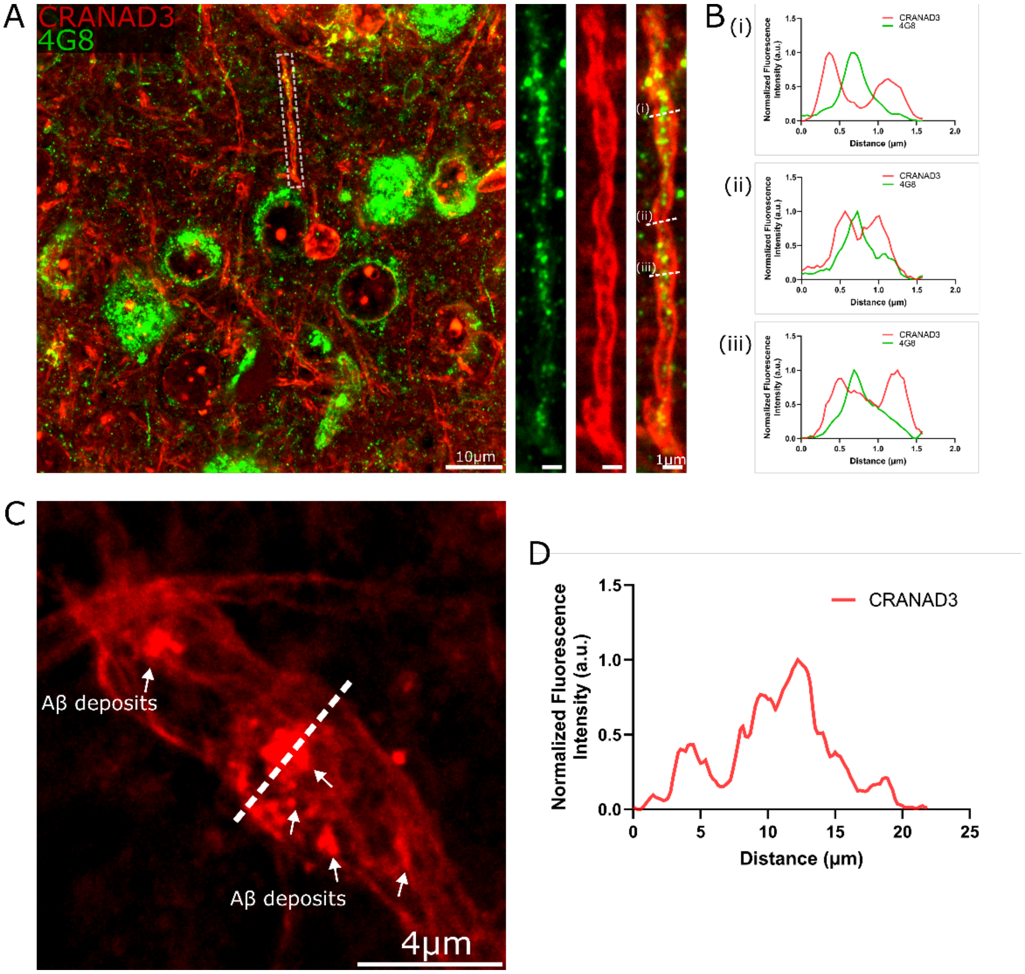


**SI Fig.7. Additional evidence of Aβ deposits transported within NLVs.**

**(A)** Non-ExM image of an AD (5xFAD) mouse brain section stained with CRANAD-3 and 4G8. Magnified views of the boxed regions (right panels) provide additional evidence that Aβ deposits can be transported within NLVs.

**(B)** Normalized fluorescence intensity profiles of CRANAD-3 and 4G8 along the dashed lines in (A).

**(C)** ExM image of a typical swollen NLV stained with CRANAD-3. White arrows indicate the location of Aβ deposits within the NLV.

**(D)** Normalized fluorescence intensity profiles of CRANAD-3 along the dashed line in (C).

Scale bars, 10 μm for left panel of (A), 1 μm for right panels of (A); 4 μm for (C).

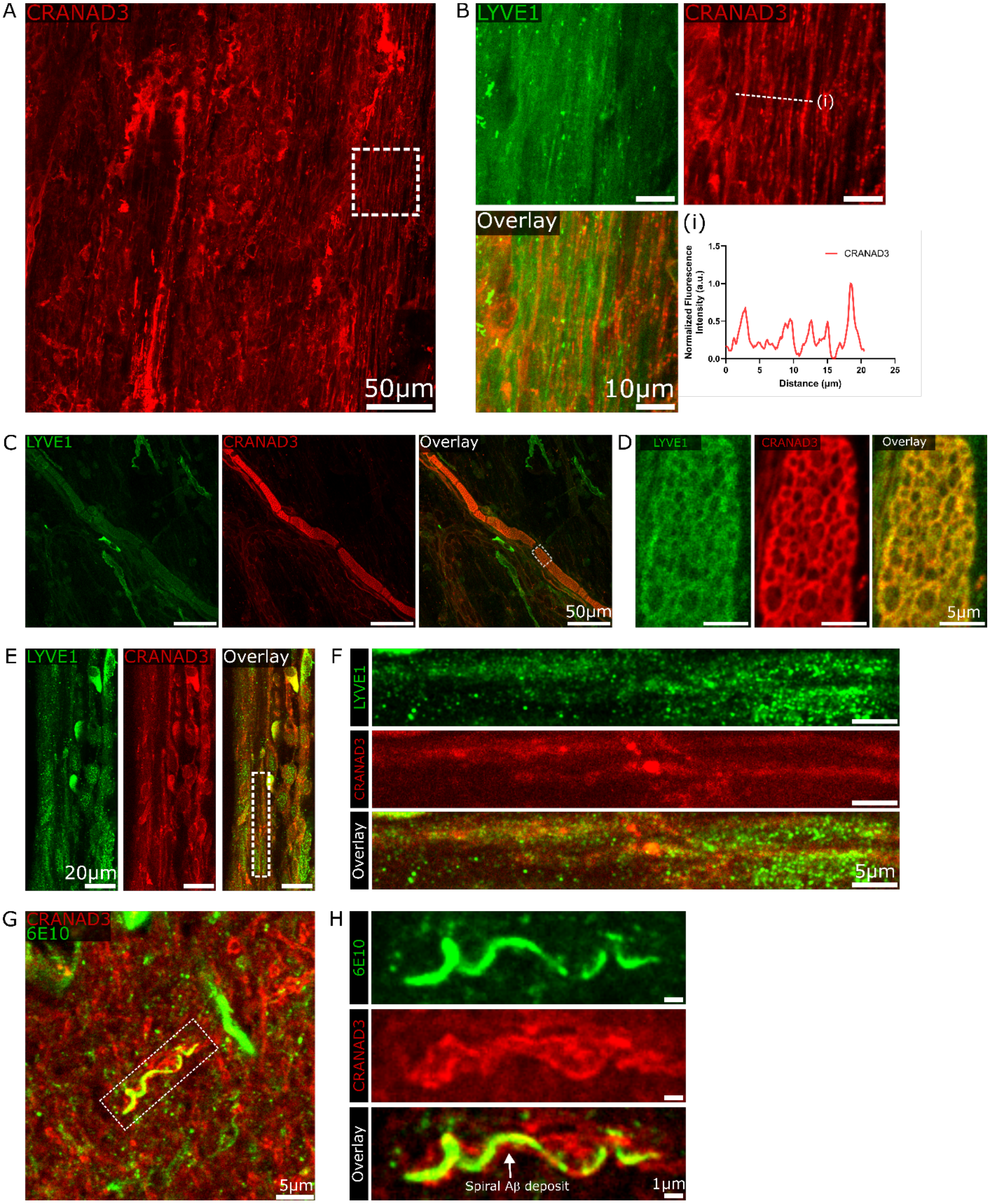


**SI Fig.8. NLVs, cobblestone-like lymphatic vessels, and lymphogenesis in mouse brain meninges.**

**(A)** Representative non-ExM image of CRANAD-3 signal in mouse brain meninges.

**(B)** Magnified view of the boxed region in (A), demonstrating the presence of NLVs within meninges. (i) Normalized fluorescence intensity of CRANAD-3 profiles along the dashed line in (B).

**(C)** Cobblestone-like lymphatic vessels in the mouse meninges.
**(D)** Zoomed-in view of the white boxed region in (C).
**(E)** Representative images illustrating lymphogenesis within the mouse brain meninges.
**(F)** Magnified view of the white boxed region in (E).
**(G-H)** Non-ExM images from mouse brain parenchyma stained with CRANAD-3 and 6E10. The magnified view of the boxed region is shown in (H), which demonstrates typical spiral Aβ deposits.

Scale bars, 50 μm for (A), 10 μm for (B); 50 μm for (C), 5 μm for (D); 20 μm for (E); 5 μm for (F); 5 μm for (G); 1 μm for (H).
